# The 2’-endo conformation of arabinose-CTP and arabinose-UTP inhibit viral polymerases by inducing long pauses

**DOI:** 10.1101/2025.08.26.672356

**Authors:** Ziyang Xiao, Arnab Das, Abha Jain, Thomas K. Anderson, Craig E. Cameron, Jamie J. Arnold, David Dulin, Robert N. Kirchdoerfer

## Abstract

Key to supporting human health in the face of evolving viruses is the development of novel antiviral drug scaffolds with the potential for broad inhibition of viral families. Nucleoside analogs are a key class of drugs that have demonstrated potential for the inhibition of several viral species. Here, we evaluate arabinose nucleotides (ara-NTP) as inhibitors of the SARS-CoV-2 and poliovirus polymerases using biochemistry, biophysics and structural biology. Ara-NTPs compete poorly with their natural counterparts for incorporation into RNA by viral polymerases. However, upon incorporation, ara-NMPs induce long polymerase pausing in both SARS-CoV-2 and poliovirus polymerase RNA elongation. Our studies suggest that subsequent nucleotide incorporation is inhibited at the catalytic step due to the 2’-endo sugar pucker of the incorporated ara-NMP.

## INTRODUCTION

The constant evolution and emergence of viruses to cause human infection represents a moving target for the design of novel antiviral drugs. Nucleotide analogs are a class of antiviral drugs that have frequently found utility in treating virus infections. These have been used to treat RNA viruses infections such as those caused by human immunodeficiency virus-1 (HIV-1)^1^, hepatitis C virus (HCV)^2^, influenza virus^3^ and coronaviruses. Nucleotide analogs have also found use in treating both infections by DNA viruses like herpesviruses as well as cancer^4,5^.

The emergence of SARS-CoV-2 in 2019 to cause COVID-19 resulted in a global pandemic and while the pandemic emergency has subsided, SARS-CoV-2 continues to circulate and evolve^6,7^. While vaccines against SARS-CoV-2 are highly effective at preventing infection or mitigating severe symptoms^8^, antiviral drugs are necessary to counter breakthrough infections and aid those unable or unwilling to be vaccinated. Nucleotide analogs typically target viral polymerases. The RNA-dependent RNA polymerase of SARS-CoV-2 is composed of a catalytic subunit, nsp12, that contains conserved polymerase motifs as well as nsp7 and nsp8 replication factors that enable the binding of RNA^9^. The nsp12 subunit is among the most conserved proteins encoded within the viral genome^10^ making it a choice target for drug discovery efforts.

Two nucleotide analog antiviral drugs have been used to treat SARS-CoV-2 infections. Remdesivir is the only FDA approved nucleotide analog for the treatment of severe COVID-19 and has been found to reduce mortality and progression to mechanical ventilation^11^. Remdesivir acts as an ATP analog and is readily incorporated into nascent RNA chains by the SARS-CoV-2 polymerase^12^. The incorporation of Remdesivir leads to delayed pausing where the polymerase continues to extend the nascent chain a further three nucleotides whereupon the 1’-cyano group of Remdesivir encounters a steric clash with nsp12 Ser861 creating translocation barrier^12,13^. Inhibition of the SARS-CoV-2 polymerase by Remdesivir can be overcome with physiological nucleotide concentrations indicating that this nucleotide analog induces polymerase pausing but not termination^14^. A second nucleotide analog, Molnupiravir, has emergency use authorization (EUA). Studies of Molnupiravir indicate that it reduces mortality and is well tolerated^15^. Molnupiravir is a cytidine analog able to base pair with both adenosine and guanosine, inducing mutations into the genome upon subsequent rounds of RNA replication. Importantly, Molnupiravir evades proofreading by the coronavirus exonuclease^16^.

To keep pace with evolving and emerging viruses and to diversify the mechanisms used to inhibit viral enzymes, it is necessary to discover and evaluate new antiviral drug scaffolds. Here we evaluate the arabinose nucleotides, ara-CTP and ara-UTP, as inhibitors of both SARS-CoV-2 and poliovirus polymerases. Arabinose nucleotides are isomers of their corresponding natural nucleotides with an inverted stereochemistry at the 2’ position of the ribose sugar. Arabinose nucleotides adopt a 2’-endo sugar pucker and are considered analogs of deoxy-cytosine triphosphate (dCTP)^17^. Ara-cytosine (ara-C), also referred to as cytosine arabinoside and cytarabine, is used for the treatment of acute myeloid leukemia and acute lymphoblastic leukemia^18^. Ara-C is phosphorylated upon entering cells to ara-CTP and competes for incorporation into DNA by DNA polymerases. DNA polymerase extension from 3’ ara-CMP is very slow, and incorporation inhibits diverse DNA polymerases including Pol-α, Pol-ε and avian myeloblastosis virus (AMV) reverse transcriptase as well as terminal transferases and DNA ligases^17,19^. Ara-thymidine (ara-T) and Ara-uridine (ara-U) have similarly been shown to inhibit DNA polymerases by competing with deoxy-thymidine triphosphate (dTTP) for incorporation into DNA by DNA Pol-β^20,21^. Like ara-C, ara-T incorporation into DNA is dependent on its conversion to ara-TTP which is less efficient using cellular thymidine kinase^21^. However, herpes simplex virus-1 and -2 (HSV-1, HSV-2) express a viral thymidine kinase that much more readily converts ara-T into its active triphosphate, providing some selectivity for targeting herpesvirus infected cells^21^. Here, we demonstrate that ara-CTP and ara-UTP are inhibitors of viral RNA polymerases *in vitro*, likely due to their 2’-endo sugar pucker and propose the development of novel antiviral nucleotide analogs from the ara-NTP scaffold.

## MATERIALS AND METHODS

### Reagent production

SARS-CoV-2 nsp7, nsp8 and nsp12 were expressed and purified as previously described^22^. Briefly nsp7 and nsp8 were expressed in *E. coli* while nsp12 was produced using the baculovirus expression system. Proteins were purified from clarified supernatants using affinity chromatography and size exclusion chromatography (**Supplementary Methods**).

Poliovirus polymerase, 3D^pol^, was expressed and purified as previously described.^23,24^ Briefly, poliovirus 3D^pol^ polymerase was expressed as N-terminal ubiquitin fusion in *E. coli*. Proteins were purified using affinity chromatography, cation exchange and size exclusion chromatography. Ubiquitin was removed during purification to create a native N terminus on poliovirus 3D^pol^ (**Supplementary Methods**).

Large RNAs for magnetic tweezers experiments were produced as previously described^25^. Briefly, the RNAs were produced as fragments and then ligated together producing a 1 kb RNA for SARS-CoV-2 polymerase experiments and a 2.8 kb double-stranded RNA for poliovirus polymerase experiments (**Supplementary Methods**).

### SARS-CoV-2 and poliovirus-catalyzed primer extensions with detection of ^32^P labeled products

Reactions to assess the incorporation of ara-NTPs by single and double nucleotide incorporations combined SARS-CoV-2 polymerase components: 0.5 µM nsp12, 1.5 µM nsp7, 1.5 µM nsp8 or 1 µM poliovirus 3D^pol^ polymerase enzyme, with 1 µM annealed synthetic RNA primer-template pairs, 100 µM ATP and 0.1 µCi/µL [α-^32^P]-ATP and were incubated at 30^°^ C. Single nucleotide incorporation assays for SARS-CoV-2 polymerase were initiated by adding either 0.1 µM CTP or UTP, or 1 mM of the corresponding nucleotide triphosphate analog (ara-CTP or ara-UTP), and allowed to proceed for 10 minutes before quenching with 50 mM EDTA. Single nucleotide incorporation assays for poliovirus polymerase were initiated by adding either 1 µM CTP or UTP, or 1 mM of the corresponding nucleotide triphosphate analog (ara-CTP or ara-UTP) and allowed to proceed for 10 minutes before quenching with 50 mM EDTA. For chain termination experiments for each polymerase, reactions were performed in the presence of the next correct nucleotide substrate (1 µM UTP or 1 µM CTP), followed by quenching with 50 mM EDTA.

To evaluate nucleotide preference of the SARS-CoV-2 polymerase complex for ara-CTP or ara-UTP, polymerase-RNA elongation complexes were assembled, and reactions were initiated with 0.1 µM CTP or UTP, along with increasing concentrations of ara-UTP or ara-CTP, respectively. These reactions were incubated for 10 minutes along with next correct nucleotide (1 µM UTP or 1 µM CTP) and then quenched with 50 mM EDTA.

Reaction products were separated by 20% denaturing urea-PAGE, analyzed by phosphorimaging and quantitated. Reactions were repeated three to four times. Data were fit by either linear or nonlinear regression using GraphPad Prism v7.03 (GraphPad Software Inc.).

### High-throughput magnetic tweezers for examining viral polymerase activity

For SARS-CoV-2 polymerase extensions, 0.1 ng of ssRNA template was bound to streptavidin-coated magnetic beads (ThermoFisher) and excess RNA was washed away. Magnetic beads bound to ssRNA templates were bound to flow cells and flushed with reaction buffer (50 mM HEPES pH 7.9, 10 mM DTT, 2 µM EDTA, and 5 mM MgCl_2_). The flow cell was then flushed with reaction buffer containing 0.6 µM SARS-CoV-2 nsp12, 1.8 µM nsp8, 1.8 µM nsp7, NTPs and ara-NTPs as indicated. Reactions were run at a constant force for 30-60 minutes. For poliovirus polymerase extensions, 0.1 ng of dsRNA template was bound to streptavidin coated magnetic beads, the beads were washed and the complexes bound to flow cells. Flow cells were rinsed in reaction buffer (50 mM HEPES pH 7.9, 5 mM MgCl_2_, 0.01% Triton X-100, 5% Superase RNase inhibitor (Life Technologies)). The flow cells were then flushed with reaction buffer containing 1.44 µM poliovirus polymerase, NTPs and ara-NTP as indicated. Reactions were run at a constant force for 60-75 minutes.

Bead displacement was converted to nucleotide progression using a linear interpolation formula. Traces were analyzed using a dwell time analysis where traces are separated into non-overlapping 10 nucleotide windows. A maximum likelihood estimation was used to analyze all the dwell times for a particular experimental condition and extract the parameters for a stochastic pausing model of polymerase extension^26^ (**Supplementary Methods**).

### Fluorescence-based RNA primer extension for incorporation and chase

To examine extension from incorporated nucleotide analogs, an RNA primer-template pair was bound to the SARS-CoV-2 polymerase and a single ara-NTP or dNTP was incorporated into the fluorescently labeled primer. The reaction as then chased with 40 µM of each of the remaining NTPs and reactions were followed over a time course to examine the formation of full-length product by denaturing urea-PAGE (**Supplementary Methods**).

### Single-particle cryo-electron microscopy for structure determination

SARS-CoV-2 nsp7, nsp8 and nsp12 were combined with an annealed RNA primer-template pair and appropriate nucleotides or nucleotide analogs were additionally added and incubated with the complex. The ara-CTP sample contained 4 mg/mL final protein concentration while the ara-UTP and UTP samples contained 8 mg/mL final protein concentrations. Samples were mixed with CHAPSO detergent just before preparing grids. Samples were spotted onto UltraAuFoil grids (Quantafoil) and plunge frozen in liquid ethane. Data was collected on a Talos Arctica (ThermoFisher) using a K3 direct electron detector (Gatan). Micrograph movies were processed in cryoSPARC^27^ to produce reconstructed density maps and coordinate models were produced using COOT^28^, PHENIX^29^ and ISOLDE^30^ using 7UOE.pdb^31^ as a starting model (**Supplementary Methods**).

### Poliovirus 3D^pol^ polymerase stopped-flow experiments

Pre-steady state nucleotide incorporation for ara-CTP, CTP, ara-UTP and UTP by the poliovirus polymerase was determined by changes in RNA template fluorescent analogs due to nucleotide incorporation into symmetrical RNA substrates and monitored by a stopped-flow apparatus. Using fluorescence traces from reaction time courses containing a range of substrate concentrations, allowed determination of the poliovirus polymerase apparent dissociation constant, K_d,app_, and polymerization rate constant, k_pol_, for each nucleotide (**Supplementary Methods**).

## RESULTS

### Incorporation of araNTP inhibits SARS-CoV-2 polymerase extension

To evaluate arabinose nucleotides (ara-NTPs) as inhibitors of viral polymerases, we tested the ability of the SARS-CoV-2 polymerase complex to incorporate and extend ara-CTP and ara-UTP. Incorporation of either ara-CTP or ara-UTP prevents further extension in a primer extension assay performed at low NTP concentrations (0.1µM NTP or 1 mM ara-NTP) (**Fig. 1A-D**). While the natural nucleotides are readily incorporated into RNA substrates, the incorporation of ara-NTP prevents further extension of the primer. This indicates an inhibition of the SARS-CoV-2 polymerase that is dependent on single nucleotide incorporation into the nascent RNA strand. We used this assay to assess the ability of each ara-NTP to compete with the corresponding natural nucleotide where extension halting at n+1 indicates incorporation of ara-NTP and extension to n+2 indicates incorporation of the natural NTP (**Fig. 1E-H**). Both ara-CTP and ara-UTP compete poorly with their corresponding natural nucleotides. The IC_50_ for ara-CTP is 30 µM in the presence of 0.1 µM CTP. Similarly, ara-UTP has an IC_50_ of 76 µM in the presence of 0.1 µM UTP. This indicates a strong preference by the SARS-CoV-2 polymerase to use natural nucleotides over their arabinose analogs.

**Figure 1:**
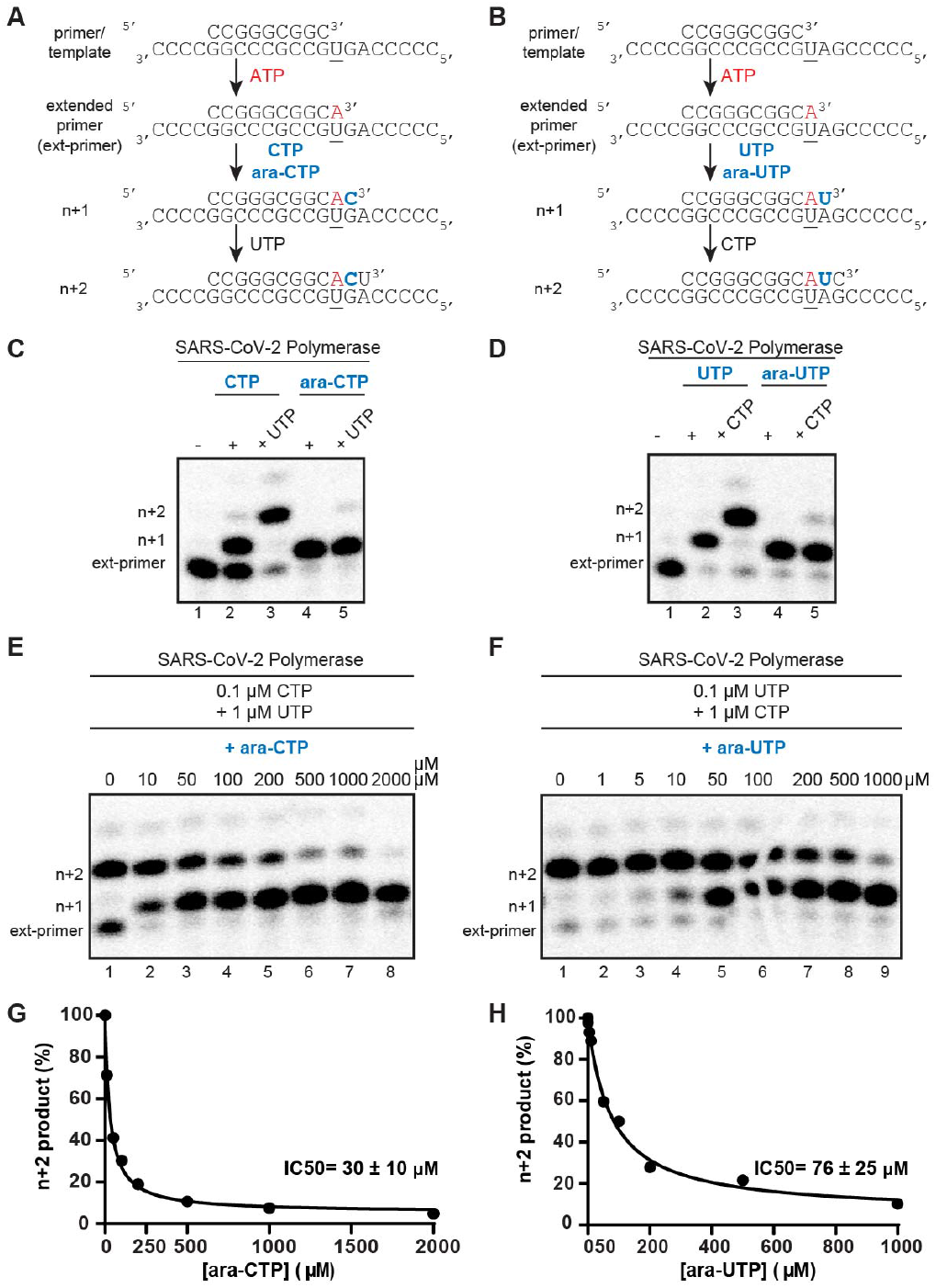
Incorporation and competition of ara-NTPs. **A**,**B)** Schematic of the primer extension assay used to evaluate SARS-CoV-2 polymerase activity. **C**,**D)** Incorporation and extension of ara-CTP and ara-UTP. SARS-CoV-2 polymerase-catalyzed incorporation of nucleoside triphosphates using CTP or ara-CTP, and UTP or ara-UTP in the absence and presence of the next correct NTP, respectively. These incorporation experiments were independently repeated at least three times with similar results. **E-H)** Competition experiment. SARS-CoV-2 polymerase-catalyzed nucleotide incorporation in the presence of increasing concentrations of ara-CTP with 0.1 µM CTP (E) and 1 µM UTP, and ara-UTP with 0.1 µM UTP and 1 µM CTP. Data was fit to a dose response curve (G and H). The IC_50_ values for ara-CTP and ara-UTP under these conditions are 30 ± 10 µM and 75 ± 25 µM, respectively.

To further investigate the mechanism of action of the ara-CTP nucleotide analog on SARS-CoV-2 RNA polymerase, we employed high-throughput magnetic tweezers (**Fig. 2, S1-2**),^14^ which take advantage of longer RNA templates, and monitor ara-NTP incorporation in the presence of all natural NTPs, including the competing one, at higher nucleotide concentrations. Here, a ∼1 kb single-strand (ss) RNA template was used to tether magnetic beads to a glass surface of a flow chamber using short, modified RNA oligonucleotides which also provides a free 3’ terminus for polymerase extension (**Fig. S1**). The polymerases elongated the primer in the presence of all NTPs with or without the nucleotide analog at an ara-CTP:CTP ratio of either 1:1 or 10:1. Polymerase extension shortens the length of the RNA tether through the conversion of the single-stranded RNA template into double-stranded (ds) RNA. By following the vertical position of the magnetic bead in real-time, we can monitor the position of the polymerase along the template with near single-base resolution and can derive the polymerase’s nucleotide addition dynamics (**Fig. 2, S2, Table S1**).

**Figure 2:**
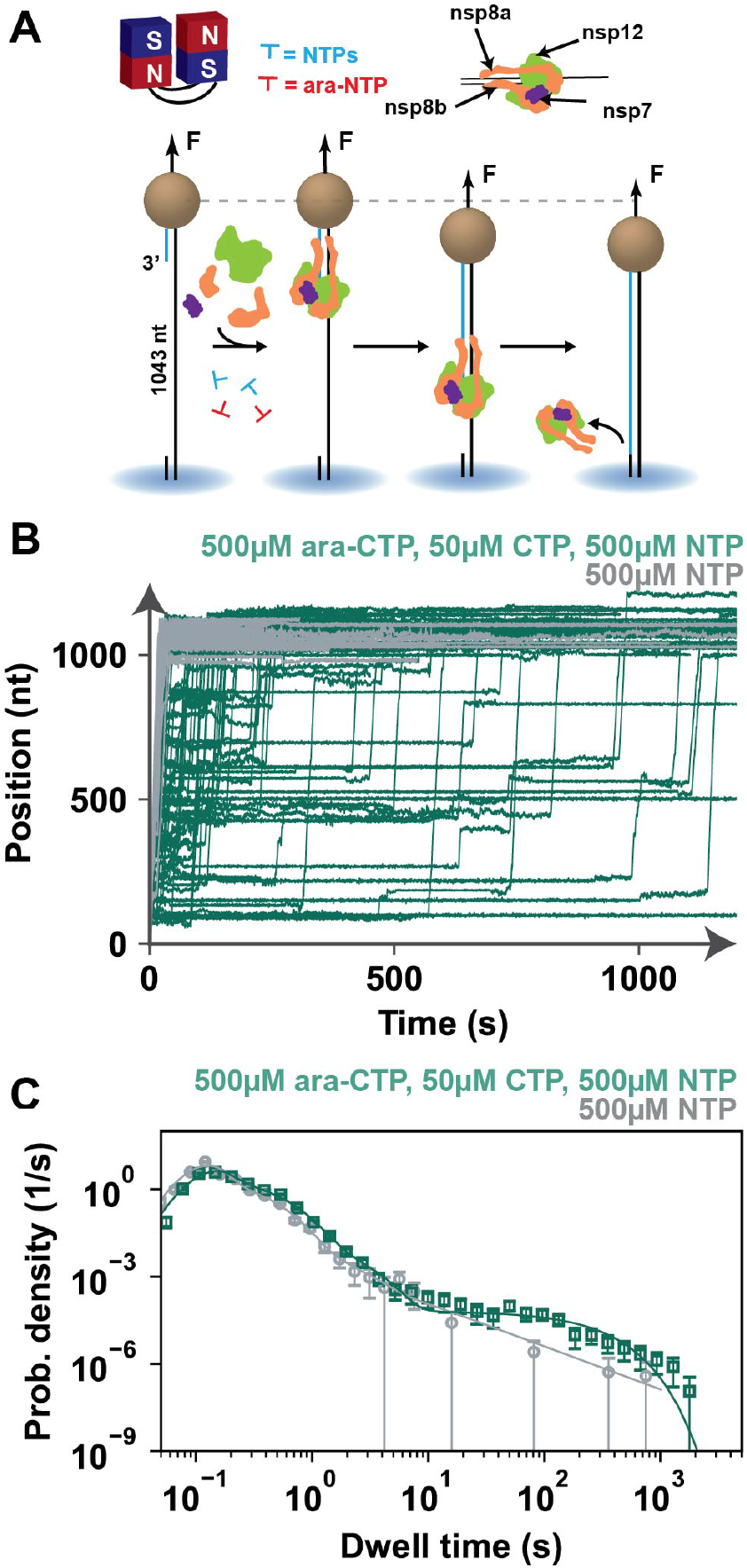
ara-CTP incorporation induces long-lived, exponentially distributed pauses in SARS-CoV-2 RNA polymerase elongation dynamics. **A)** Schematic of the magnetic tweezers assay to monitor SARS-CoV-2 polymerase RNA synthesis activity. **B)** SARS-CoV-2 polymerase RNA synthesis activity traces for either 0 µM ara-CTP and 500 µM NTP (grey), or 500 µM ara-CTP, 50 µM CTP and 500 µM other NTP (dark green). **C)** Dwell time distributions extracted from the SARS-CoV-2 polymerase activity traces described with the same color code as indicated in A). The solid lines are maximum likelihood estimation (MLE) fits of the pause-stochastic model. The error bars are one standard deviation from 1,000 bootstraps of the data sets.

Previous studies of the SARS-CoV-2 RNA polymerase using magnetic tweezers have revealed rapid incorporation of nucleotides with evidence of distinct on-pathway polymerase states associated with slower nucleotide incorporation states as well as an off-pathway long-lived kinetic state assigned to backtracked polymerases^32^. The slow nucleotide addition pathway has been suggested to represent a lower fidelity polymerase state (Pause 1) and the very slow nucleotide addition pathway as polymerase extension from the products of nucleotide misincorporations (Pause 2)^26^.

In the presence of ara-CTP, we monitored an increase in the number of long-lived pauses interrupting the activity traces (**Fig. 2, S2, Table S1**), increasing the average RNA replication time while not affecting the mean product length. The unaffected product length in the presence of ara-CTP at higher NTP concentrations suggests that contrary to bulk biochemistry experiments above, ara-CTP incorporation by the SARS-CoV-2 RNA polymerase introduces strong polymerase pausing but does not act as a terminator.

We performed a dwell time analysis of the magnetic tweezers polymerase elongation traces, scanning traces with a non-overlapping 10 nt window to measure the amount of time polymerase took to incorporate ten successive nucleotides (dwell times). The dwell time duration is the temporal signature of the biochemical process that dominated the 10 nt addition. The resulting dwell times are assembled into a probability density distribution represented in a log-binned histogram (**Fig. 2, S2**)^26^. These distributions were then fitted using a stochastic pausing translocation model from which we derive the nucleotide addition rate, probabilities for polymerase entering a slow or very slow nucleotide addition pathway and backtracked states, as well as the exit rate of each nucleotide addition pathway^26,32^ (**Table S1**).

Both the slow and very slow nucleotide addition pathways (Pause 1 and Pause 2) are exponentially distributed. In the absence of ara-CTP, the long-lived pause is best described by a power law in the absence of ara-CTP as previously reported for SARS-CoV-2 polymerase^14^. In the presence of ara-CTP, the long-lived pause is best described by an exponential distribution, which we term the ara-CTP pause. The exponential distribution of the ara-CTP pause indicates that such a pause is exited through a single rate-limiting kinetic step with an exit rate k_ara-CTP_=0.005 s^-1^, i.e. the ara-CTP pause has an average lifetime of ∼200 s (**Table S1**). The nucleotide addition, slow and very slow nucleotide addition pathway’s exit rates were largely unaffected by the addition of ara-CTP as compared to the SARS-CoV-2 RNA polymerase with 500 µM NTPs. The slight difference in observed in Pause 2 exit rate at the highest stoichiometry are likely due to the difficulty in fitting the Pause 2 exponential when the ara-CTP pause becomes dominant in the dwell time distribution.

The strong pausing observed in the magnetic tweezers for ara-CTP and SARS-CoV-2 could be recapitulated in bulk biochemistry assays. In these assays, ara-CTP or ara-UTP is incorporated into primer-template pairs by the SARS-CoV-2 polymerase and then chased with 40 µM of each of the remaining NTPs. Over the course of the experiment, incorporated ara-NMPs are extended to a full-length product demonstrating escape from the strong polymerase pausing (**Fig. S3**). Despite SARS-CoV-2 polymerase’s ability to extend from both ara-CMP or ara-UMP incorporations, ara-CMP proved more difficult to extend from with significant inhibited product remaining after an hour whereas, ara-UMP could be extended to full-length product in 30 min. The lack of significant stalling beyond the incorporated ara-NMP supports the finding from the bulk biochemistry experiments above in that a single ara-NMP incorporation is sufficient to inhibit polymerase and differs from nucleoside analogs such as Remdesivir which pause the SARS-CoV-2 polymerase after several additional nucleotide incorporations^12^.

### Ara-NTPs bind the SARS-CoV-2 polymerase active site and are readily incorporated

To further examine the mechanism of action of ara-NTPs against SARS-CoV-2, we used cryo-electron microscopy (cryo-EM) to determine several structures with incorporated ara-NMPs (**Fig. 3, S4-7, Table S2**). The 3.3 Å structure of the SARS-CoV-2 polymerase complex with incorporated ara-CMP shows good resolution for the nsp12 catalytic subunit as well as nsp7 and two nsp8 co-factors. We observe 30 base pairs of dsRNA and six single-stranded nucleotides of downstream template. The polymerase has incorporated one ara-CMP and has translocated to allow the binding of an incoming ara-CTP (**Fig. 3A, S4, S8**). We also observe an ara-CTP bound to the N-terminal nidovirus RNA polymerase associated nucleotidyltransferase (NiRAN) domain in a base-in conformation (7ED5^33^ and 7UOB^31^) with H-bonds of Cys53 to the arabinose 2’OH and Tyr257 to the cytidine base. The binding of ara-CTP to the NiRAN appears to take advantage of the catalytic metals in the NiRAN active site as has been observed in several in-facing and out-facing NTPs (6XEZ^34^ and 7CYQ^35^). We determined a similar structure with an incorporated ara-UMP though without reliable density to model an incoming ara-UTP or NiRAN bound nucleotide (**Fig. 3B, S5, S8**). The ara-nucleotides were modeled with 2’-endo sugar puckers as has been suggested by previous structural studies^36,37^ and supported by the reconstructed maps. In both incorporated ara-NMP structures, the nucleotide analogs participate in Watson-Crick base pairing interactions with the corresponding template base and the RNAs are in post-translocated states.

**Figure 3:**
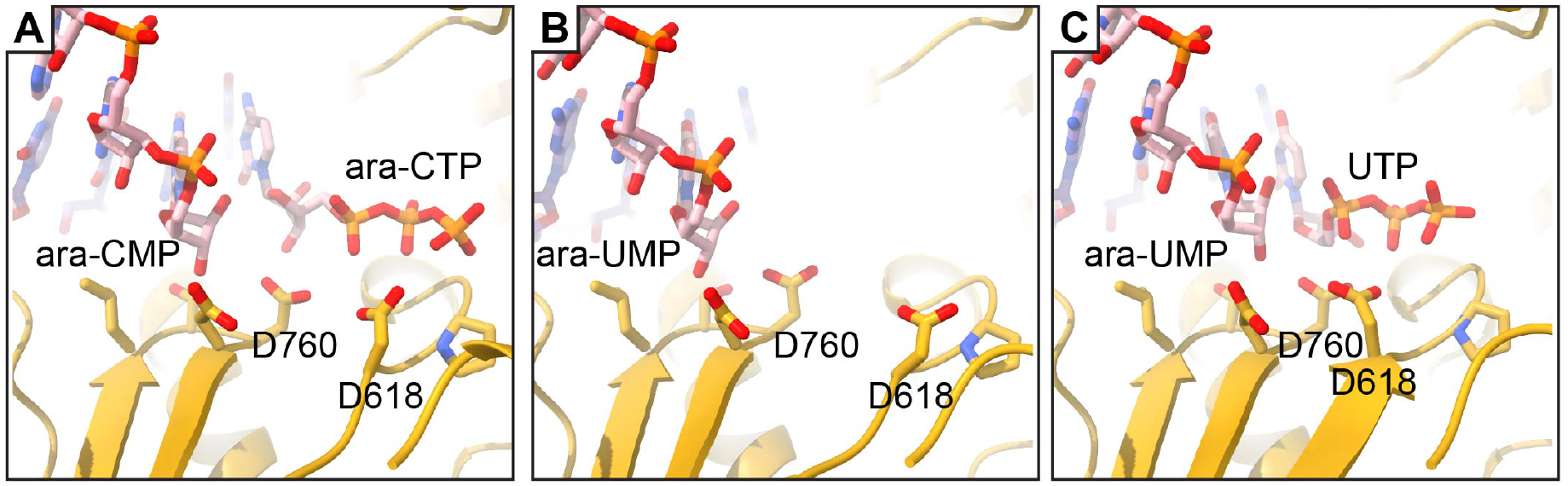
SARS-CoV-2 polymerase active site poses for incorporated ara-NMPs. **A)** Coordinate model of the polymerase active site with incorporated ara-CMP with incoming ara-CTP. **B)** Coordinate model of the polymerase active site with incorporated ara-UMP. **C)** Coordinate model of the polymerase active site with incorporated ara-UMP with incoming UTP. Nucleotide analogs are labeled. D760 corresponds to a conserved polymerase motif C. D618 is a conserved residue of polymerase motif A which shifts position to close the RNA polymerase active site when a nucleotide is bound^38^.

Because the biochemical studies above showed that ara-NMP incorporation induced polymerase pausing and not termination, we also determined a structure with an incorporated ara-UMP and an incoming UTP (**Fig. 3C, S6, S8**). As a control, we also include a structure of polymerase that has incorporated two UMP (**Fig. S7, S8**).

### Structural basis for poor SARS-CoV-2 polymerase ara-NTP selectivity over natural nucleotides

In both RNA and DNA polymerases, incoming nucleotides are recognized by a network of H-bonding and/or hydrophobic interactions with the 2’ and 3’ substituents of the ribose or deoxyribose sugar^1,38–40^. These 2’ and 3’ substituents also influence the preferred pucker of the sugar where RNA bases favor a 3’-endo conformation and DNA bases prefer a 2’-endo conformation^41,42^. As the SARS-CoV-2 polymerase structure with incorporated ara-CMP also contains an incoming ara-CTP, we compared the incoming ara-CTP pose to SARS-CoV-2 polymerase structures with natural incoming nucleotides such as our araUMP with incoming UTP structure or the published structures of 3’-deoxynucleotide terminated RNA with incoming CTP or UTP^31^. In the ara-CMP incorporated structure, the arabinose sugar of the incoming ara-CTP is poorly resolved with stronger density for base and triphosphate. The weaker density of the arabinose sugar than adjacent base and phosphate moieties implies a higher flexibility for the sugar of the incoming ara-CTP. The incoming ara-CTP base makes Watson-Crick base pairing interactions with the templating base. The ara-CTP triphosphate has an altered position compared to incoming CTP (7UOE.pdb^31^) (**Fig. S9**) resulting in an altered coordination of the polymerase catalytic site magnesium ions. In addition, despite the binding of an incoming nucleotide, the ara-CTP polymerase structure remains in an open polymerase state. It has previously been hypothesized that engagement with incoming nucleotide 2’ and 3’ hydroxyl groups results in active site closure prior to catalysis^38^. As the ara-UMP incorporated with incoming UTP structure adopts a closed pre-catalytic polymerase, we assign the failure to close the active site for catalysis to the incoming ara-CTP rather than the incorporated ara-CMP. Together, the lack of stabilization for the binding of the arabinose sugar, an altered binding of the triphosphate to the polymerase catalytic center and the failure to close the polymerase active site all likely contribute to poor competition of ara-NTPs with natural NTPs for incorporation into the growing RNA strand.

### Structural mechanism of action for ara-NMP induced polymerase pausing

Despite not competing well with natural nucleotides, incorporation of ara-NMPs into nascent chains leads to strong polymerase pausing as described above in magnetic tweezers and bulk biochemistry experiments. A comparison of the ara-UMP and ara-CMP incorporated SARS-CoV-2 polymerase structures with published structures shows no large differences in either the protein or RNA components (**Fig. S9**). Polymerases of positive-sense RNA polymerases undergo a multi-step cycle to incorporate a nucleotide into nascent RNA. These include NTP binding, active-site closure, catalysis, active-site opening and translocation^38,43^. Each of these steps could potentially be inhibited by nucleotide analog polymerase inhibitors. The structures with incorporated ara-CMP and ara-UMP have each adopted post-translocated states suggesting that for each nucleotide analog, the post-translocated state is favored at equilibrium and translocation is not strongly inhibited by ara-NMP incorporation. Moreover, as we showed above, single incorporation of ara-NMPs is sufficient for polymerase inhibition and incorporation and extension data suggest no additional strong pausing by the SARS-CoV-2 polymerase ruling out a delayed pause. Upon resumed extension from the long ara-NMP pauses, nucleotide addition is rapid suggesting that the long ara-NTP pauses are the result of an incorporated ara-NMP that interferes with subsequent nucleotide addition. The polymerase active site conformation with incorporated ara-NMP does not prevent NTP binding as evidenced by the ara-UMP incorporated structure with incoming UTP. Moreover, the binding of the natural UTP in the structure promotes active site closure identically to other SARS-CoV-2 structures with the expected shifting of polymerase catalytic motifs A and C to close the active site and UTP positioning identically to incoming natural nucleotides^31^ (7UO9.pdb) (**Fig. S9**).

Close examination of the incorporated ara-NMPs in each structure shows an increased distance and an angling away of the 3’ hydroxyl group from the polymerase active site and the α-phosphate of an incoming NTP where present (**Fig. 4**). This shift is due to the 2’-endo sugar pucker of the arabinose sugar which alters the 3’ hydroxyl position.

**Figure 4:**
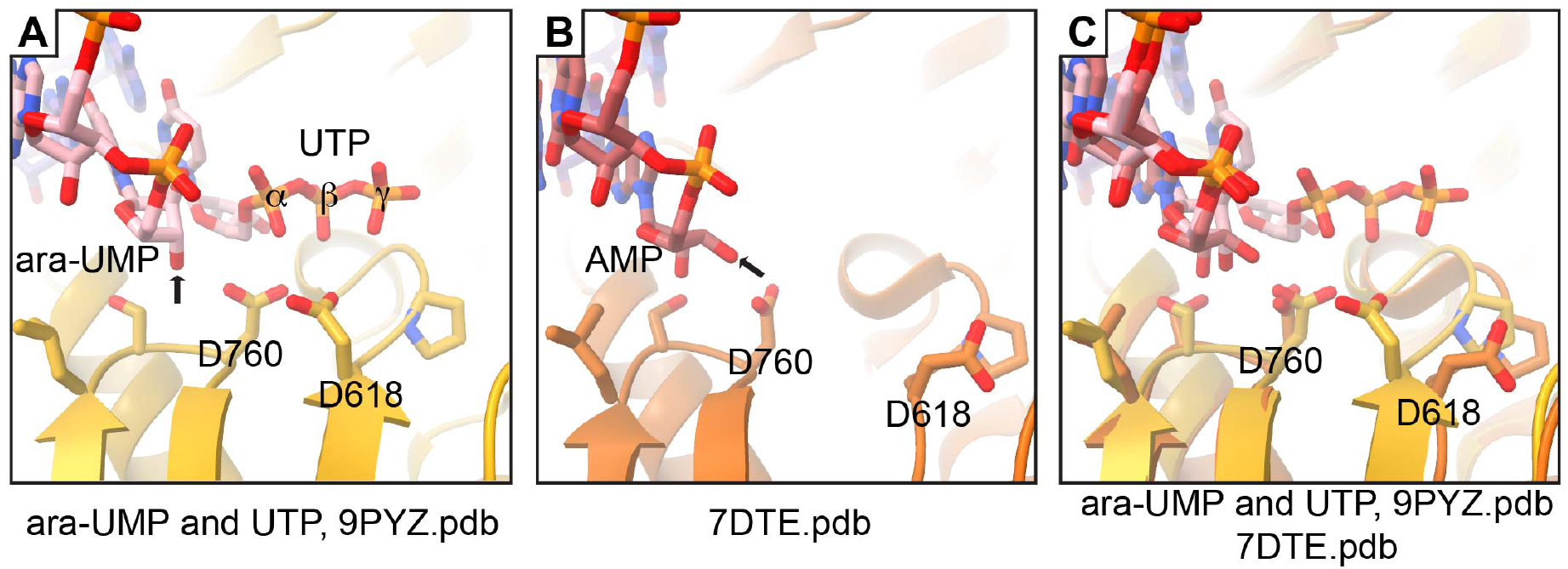
The 2’-endo sugar pucker of ara-NMPs reorients the 3’ hydroxyl for inefficient nucleophilic attack. **A)** The SARS-CoV-2 polymerase with incorporated ara-UMP and incoming UTP. The 3’ hydroxyl of the arabinose sugar in the 2’-endo configuration is indicated with an arrow. The phosphates of the incoming UTP are labeled. **B)** A previously determined polymerase structure^51^ with a 3’ AMP. The 3’ hydroxyl of the ribose sugar in the 3’-endo configuration is indicated with an arrow. **C)** Superposition of the structures in A) and B) comparing different orientations of the 3’ hydroxyl. D760 and D618 indicate conserved polymerase motifs C and A respectively. See also **Fig. S9**.

Using the incorporation and chase assay described above for ara-NTPs, we used 2’ deoxynucleotides (dNTPs) which also have a 2’-endo sugar pucker. While both dCTP and dUTP are incorporated into synthetic RNAs by the SARS-CoV-2 polymerase, further extension of the primer using the remaining NTPs indicates a strong pausing of polymerase extension similar to that observed for the ara-CTP and ara-UTP (**Fig. S10**).

### Ara-CTP and ara-UTP inhibition of poliovirus polymerase

To evaluate the extent to which arabinose nucleotides may broadly inhibit viral RNA polymerases, we tested the effects of both ara-CTP or ara-UTP against poliovirus RNA polymerase. (**Fig. S11**). The IC_50_ for both ara-CTP and ara-UTP are 1000 µM in the presence of 1 µM CTP or UTP. To further elucidate the efficiency of incorporation we determined the kinetic parameters: the *k*_pol_ value, the maximal rate constant for nucleotide incorporation, and the *K*_d,app_ value, the apparent dissociation constant for NTP binding for each ara-NTP and the corresponding natural nucleotide using a stopped-flow fluorescence apparatus^44,45^. The differences in efficiency between ara-CTP and ara-UTP and natural nucleotides are 50- and 90-fold lower respectively (**Table S3-4**).

We also assessed the action of ara-UTP and ara-CTP using our high-throughput magnetic tweezers assay. Differing from our magnetic tweezers assay for SARS-CoV-2 polymerase, the assay to assess poliovirus polymerase uses a dsRNA template with a short 3’ hairpin. Polymerase extension from the 3’ hairpin converts the dsRNA tether into ssRNA, increasing the length of the tether and shifting the position of the magnetic bead. The change in magnetic bead vertical position is then converted to nucleotides extended^46^ **(Fig. S13-14**). Inclusion of ara-CTP or ara-UTP at ratios of 1:10 or 1:5 to their natural NTPs had very mild effects on pause rates or probabilities. Increasing the ratios to 1:1, we observe a 3-fold increase in ara-NTP pause probabilities for each analog (**Tables S5-6**).

## DISCUSSION

Ara-CTP and ara-UTP induce strong pausing and inhibition of SARS-CoV-2 and poliovirus polymerases. We have shown that single ara-NTP incorporation is sufficient to induce long-lived pausing of SARS-CoV-2 polymerases. However, we observe a poor ability of ara-NTPs to compete with their corresponding natural NTPs likely owing to the ara-NTPs sub-optimal binding to polymerase active sites. The sub-optimal binding of the ara-NTPs may be due to altered base-pairing or base stacking^36,37^, poor coordination of the arabinose sugar all potentially leading to a failure of the polymerase active site to adopt a closed pre-catalytic configuration. We also observed an altered engagement of polymerase active site by the ara-CTP triphosphate compared to CTP (7UOE.pdb^31^) though this interaction may represent the initial binding of NTPs to the open polymerase state^38,47^.The pausing caused by the incorporation of ara-NMPs was not the result of poor RNA translocation, subsequent NTP binding or active site closure, nor is the polymerase active site distorted to prevent catalysis. Previous work examining the potential of ara-UTP as an inhibitor of SARS-CoV-2 polymerase similarly found that ara-UMP incorporation does not result in termination and that ara-UTP is not a preferred substrate over natural UTP. However, the longer endpoint assays adopted in this study did not observe the prolonged pausing induced by ara-UMP incorporation or inhibition of SARS-CoV-2 polymerase primer extension^48^. The immediate pausing caused by ara-NMP incorporation differs in its mechanism from the nucleotide analog Remdesivir which introduces a translocation barrier and strong pausing after the incorporation of an additional three nucleotides due to a steric clash in the polymerase dsRNA exit channel^12–14^.

We propose that ara-CTP and ara-UTP inhibit viral polymerases by inducing a strong pause immediately after single nucleotide analog incorporation and translocation, driven by the 2’-endo sugar pucker conformation being non-conducive for nucleophilic attack on the α-phosphate of incoming NTPs. Ara-CTP has long been used as a dCTP analog for the treatment of acute myeloid leukemia and also inhibits DNA polymerases^17^ at a catalytic step^49^ despite ara-CMP adopting a 2’-endo conformation similar to dCMP^36,37^. The inhibition of DNA polymerases may be due to the higher pseudo-rotation phase angle of ara-CMP than dCTP resulting in an increased sugar pucker^36^ and altered active site engagement. It has also been found that in contrast to dCMP, ara-CMP does not fluctuate between 2’-endo and 3’-endo conformers^36,37^. Both SARS-CoV-2 and poliovirus polymerases show similar long-lived pauses induced upon ara-CMP and ara-UMP incorporation. These data suggest that the adoption of strong 2’-endo sugar puckers may be a common mechanism of inhibition across viral polymerases and that nucleoside scaffolds with 2’-endo sugars may find use in the development of broad-spectrum RNA virus antivirals.

While the use of ara-nucleosides is an attractive scaffold for further discovery, significant improvements in the ara-NTPs studied here will be necessary to bypass several shortcomings in these nucleotide analogs prior to therapeutic use. The steric clash between the ara-NTP 2’OH and the H6 pyrimidine base hydrogen^36,37^ may result in weaker base pairing with the template base and reduce base stacking leading to weakened binding of ara-NTPs to SARS-CoV-2 nsp12 NTP-binding site. 2’fluoro-ara-NTPs which replace the 2’OH with a 2’F should be evaluated against viral RNA polymerases as these analogs make pseudo-H-bonds to their own H6 rather than creating a steric clash^36,37^. This would pre-organize the nucleotide base for base pairing with template nucleotides which may partly overcome the observed issue with competition of ara-NTPs with natural NTP substrates. For reasons which remain unclear, the ara-CTP appears to compete better with CTP than ara-UTP competes with UTP *in vitro* and intracellular CTP levels are typically much lower than other nucleotides^50^. We therefore suggest the development of ara-CTP as a better scaffold for elaboration than ara-UTP. There are also cytotoxicity concerns regarding the use of ara-NTPs as antiviral drug scaffolds because ara-cytidine, the nucleoside of ara-CTP, is used clinically to treat leukemia. Ara-cytidine is converted into ara-CTP in cells and exerts its anti-cancer effects by competing with dCTP for incorporation into DNA with Pol-α and Pol-ε^17,19^. Like its effect on viral polymerases, incorporated ara-CMP is extended poorly by DNA polymerases leading to replication stress and cell death through the stalling of replication forks and the creation of double-stranded DNA breaks^19^. The development of nucleotide analogs that target the SARS-CoV-2 replication machinery has been difficult due to the proofreading activity of the viral exonuclease. While we do not evaluate the efficacy of ara-nucleosides in reducing SARS-CoV-2 titers and do not test the ability of the viral exonuclease to remove ara-NTPs from RNA termini, proofreading of incorporated ara-NMPs has been examined in several DNA polymerases and the ara-NTP analogs are efficiently excised by those polymerases possessing 3’ to 5’ exonuclease activity^17,19^. The excision of the nucleotide from nascent RNA in SARS-CoV-2 replication would be expected to narrow the therapeutic window in which these analogs could be used as drugs.

In summary, we present ara-NTPs as a scaffold for further elaboration for the development of novel antiviral drugs for targeting diverse RNA viruses. We propose that the adoption of a 2’-endo sugar pucker in incorporated ara-NMPs induces long pausing in viral polymerases and identify several limitations of the current ara-NTPs that may be overcome with additional medicinal chemistry efforts. The introduction of promising scaffolds for viral polymerase inhibition and preliminary characterization of those scaffolds’ mechanisms of inhibition will be essential to address evolving viral threats both from existing and emerging viruses.

## ACKNOWLEDGMENTS

Some of this work was performed in the Cryo-EM Research Center (CEMRC) in the Department of Biochemistry at the University of Wisconsin-Madison. This work was supported by funding NIH/NIAID AI158463 to R.N.K.; NIH/NIAID AI161841 to C.E.C. and J.J.A. D.D. was funded by BaSyC – Building a Synthetic Cell” Gravitation grant (024.003.019) of the Netherlands Ministry of Education, Culture and Science (OCW) and the Netherlands Organisation for Scientific Research (NWO), and NWO funding OCENW.XL21.XL21.115.

## DATA AVAILABILITY

Structure coordinate data and reconstructed maps are available for download at the Protein Data Bank (PDB) or the Electron Microscopy Data Bank (EMDB). The ara-CMP and ara-CTP structure has accession numbers pdb_00009BLF, EMD-44654. The ara-UMP structure has accession numbers pdb_00009PYW, EMD-72038. The ara-UMP and UTP structure has accession numbers pdb_00009PYZ, EMD-72053. The structure with two incorporated UMP has accession numbers pdb_00009PZ0, EMD-72054.

