## Supplemental tables and figures for "The 2’-endo conformation of arabinose-CTP and arabinose-UTP inhibit viral polymerases by inducing long pauses"

**Supplemental Data:**
Supplemental Material and Methods
Supplemental Tables S1-4
Supplemental Figures S1-12
Supplemental References

### SUPPLEMENTAL MATERIALS AND METHODS

#### Reagent production

Expression and purification of SARS-CoV-2 polymerase components

All SARS-CoV-2 nsps were codon optimized and synthesized from Genscript. Protein sequences correspond to GenBank UHD90671.1. SARS-CoV-2 nsp7 and nsp8 genes were cloned into pET46 plasmids with N-terminal 6xHis tags and TEV protease sites. The SARS-CoV-2 nsp12 (RNA dependent RNA polymerase) gene was cloned into the pFastBac plasmid with C-terminal TEV protease site and double Strep tags.

Recombinant SARS-CoV-2 nsp7 and nsp8 were individually expressed in Rosetta2 pLysS *E. coli* cells (Novagen). Cell cultures were grown at 37°C until they reached $OD_{600}$ of 0.6-0.8, then induced by the addition of isopropyl β-d-1-thiogalactopyranoside (IPTG) to a final concentration of 500 µM and incubated at 16°C for 16 hours. Cells were pelleted and resuspended in wash buffer (25 mM HEPES pH 7.4, 300 mM sodium chloride, 30 mM imidazole and 2 mM dithiothreitol (DTT)) and lysed using a Microfluidizer (Microfluidics). Lysates were cleared by centrifugation and filtration with 0.45 µm filters. The protein was purified using Ni-NTA agarose beads (Qiagen), washing with wash buffer and eluting same buffer containing 300 mM imidazole. The eluted protein was dialyzed at 4°C overnight into Dialysis buffer (25 mM HEPES pH 7.4, 300mM sodium chloride, and 2 mM DTT) together with 1% (w/w) TEV protease. The digested protein was then passed over Ni-NTA beads column again to remove the uncleaved proteins. Digested proteins were further purified on Superdex 200 Increase 10/300 GL column (Cytiva) in 25 mM HEPES pH 7.4, 300 mM sodium chloride and 2 mM DTT. Fractions containing the protein of interest were concentrated, aliquoted, flash frozen with liquid nitrogen and stored at -80°C.

The pFastBac plasmid carrying SARS-CoV-2 nsp12 was transformed into DH10Bac *E. coli* cells (Life Technologies) to produce recombinant bacmids. The bacmids were then transfected into Sf9 cells using Cellfectin II (Life Technologies). Recombinant baculovirus was amplified twice in the Sf9 cells. P2 stocks of baculovirus were used to infect Sf21 cells for protein expression and expression was allowed to proceed for 36 hours. Infected cells were pelleted by centrifugation and resuspended in 25 mM HEPES pH 7.4, 300 mM sodium chloride, 1 mM magnesium chloride, and 5 mM DTT. 143 µL BioLock (IBA) was added per liter of cell culture before lysis in a Microfluidizer (Microfluidics). Lysates were cleared by centrifugation and filtration through 0.45 µm filters. Strep-tagged nsp12 was collected on Streptactin superflow agarose (IBA), washed with lysis buffer and then eluted with the same buffer containing 2.5 mM desthiobiotin. Nsp12 was further purified on Superdex 200 Increase 10/300 GL column (Cytiva) in 25 mM HEPES pH 7.4, 300 mM sodium chloride, 100 µM magnesium chloride, and 2 mM Tris(2-carboxyethyl)phosphine (TCEP). Fractions of nsp12 were concentrated, flash frozen in liquid nitrogen and stored at -80°C.

Expression and purification of poliovirus polymerase

Poliovirus polymerase, 3Dpol, was expressed and purified as previously described.[1,2] Briefly, expression was performed at 25 °C by auto-induction, cells harvested, lysed by French Press, subjected to PEI precipitation followed by ammonium sulfate precipitation, Ni-NTA chromatography, cleavage by Ulp1, phosphocellulose chromatography, and size exclusion chromatography. The protein was concentrated using Vivaspin concentrators.

Production of ssRNA for magnetic tweezers experiments

ssRNA for examining the extension activity of the SARS-CoV-2 polymerase was produced as several RNA fragments that were then annealed and ligated together[3]. To obtain the different parts of the RNA construct, pBB10 and pLM659 template DNA fragments[4,5] were amplified via PCR and purified (Monarch PCR and DNA cleanup kit, DNA primers from Biomers.net or Integrated DNA Technology (IDT)). DNA products were *in vitro* transcribed (NEB HiScribe T7 High Yield RNA Synthesis Kit). Labeled RNAs were created by including Biotin-16-UTP or digoxygenin-11-UTP in appropriate *in vitro* transcription reactions. RNA transcripts that required ligation to the next fragment at the 5' end were treated with RNA 5' Polyphosphatase (Biosearch Technologies). Seven RNA fragments were annealed and ligated with T4 RNA ligase 2 (NEB) to assemble the final RNA. The final RNA structure is a 499 bp double-stranded RNA stem with a 20 nt loop flanked by dsRNA regions. The strand containing the stem loop contains a 343 nt digoxygenin-labeled RNA at its 5' end and a 443 nt biotin-labeled RNA at its 3' end. Each of these labeled RNAs lie in the dsRNA flanking regions. Upon application of >21 pN of force, the hairpin melts to present ssRNA as a polymerase template[6].

Sequence of the single-stranded RNA template (open hairpin; 5' → 3', 1043 nt):

GUUCUACAUAGCGUGCAGACGUGAAUUUAAUCUCGCUGACGUGUAGACACAGUGCGUCUGCUGUCGG
GUCCCUCUGGUGACUGGGUAGUUGGACUUGCCCUUGGAAGACAUAGCAAGACCCUGCCUCUCUAUUG
AUGUCACGGCGAAUGUCGGGGAGACAGCAGCGGCUGCAGACAUCAGAUCGGAGUAAUACUCUCCGUA
ACUGGCCUUCUCUGAAUUCCGACGUUGUUAAGAUGGCAGAGCCCGGUAAUCGCUACUUGACCAGAUAA
GCUUUCCGUGGAUGGUUUAGAGGAAUCACAUCCAAGACUGGCUAAGCACGAAGCAACUCUUGAGUGUA
AAAUUGUUGUCUCCUGUAUUCGGGAUGCGGGUACUAGAUGACUGCAGGGACUCCGACGUUAAGUACA
UUACCCCGUCAUAGGCGCCGUUCAGGAUCACGUUACCGCCAUAAGAUGGGAGCAUGACUUCUUCUCC
GCUGCGCCCACGGAUCCAGUAGUGAUUAACAUUCGACAGCAUGCGCACUAAUCACUACUGGAUCCGUG
GGCGCAGCGGAGAAGAAGUCAUGCUCCCAUCUUAUGGCGGUAACGUGAUCCUGAACGGCGCCUAUGA
CGGGGUAAUGUACUUAACGUCGGAGUCCCUGCAGUCAUCUAGUACCCGCAUCCCGAAUACAGGAGACA
ACAAUUUUACACUCAAGAGUUGCUUCGUGCUUAGCCAGUCUUGGAUGUGAUUCCUCUAAACCAUCCAC
GGAAAGCUUAUCUGGUCAAGUAGCGAUUACCGGGCUCUGCCAUCUUAACAACGUCGGAAUUCAGAGAA
GGCCAGUUACGGAGAGUAUUACUCCGAUCUGAUGUCUGCAGCCGCUGCUGUCUCCCCGACAUUCGCC
GUGACAUCAAUAGAGAGGCAGGGUCUUGCUAUGUCUUCCAAGGGCAAGUCCAACUACCCAGUCACCAG
AGGGACCCGACAGCAGACGCACUGUGUCUACACGUCAGCGAGAUUAAAUUCACGUCUGCACGCUAUGU
AGAACCCUCAGCCAACUCGGUCGCGUCGGA

Production of dsRNA templates for magnetic tweezers experiments

The construct employed here, which has been previously described in detail in Quack et al.[7] is made of a 4 kb long single-stranded RNA tether to which four ssRNAs are annealed: a biotin-labeled strand to attach to the streptavidin-coated magnetic bead, a spacer, ~2.9 kb template, and a digoxygenin-labeled strand to attach the magnetic bead to the surface glass surface. The template RNA has a small hairpin with the sequence ACGCUUUCGCGT followed by 15 U residues to initiate poliovirus polymerase catalyzed RNA synthesis via primer extension.

Annealing RNA for primer extensions with detection of $^{32}$P labeled products

For nucleotide incorporation assays, dsRNA substrates were prepared by annealing a 5′-CCGGGCGGC RNA primer with either a 5′-CCCCCAGUGCCGCCCGGCCCC RNA template for CTP or ara-CTP incorporation, or a 5′-CCCCCGAUGCCGCCCGGCCCC RNA template for UTP or ara-UTP incorporation. dsRNA substrates were produced by annealing 10 μM RNA oligonucleotides in 10 mM Tris pH 8.0, 1 mM EDTA and 50 mM NaCl in a Progene Thermocycler (Techne). Annealing reaction mixtures were heated to 90°C for 1 min and slowly cooled (5°C/min) to 10°C.

Annealing RNA for primer extensions with detection of 6-FAM labeled products and cryo-EM

Pair 1 (ara-CTP and dCTP extension assay):
Primer RNA: CAUUCUCCUAAGAAGCUAUUAAAAUCACA
Template RNA: AAAAACCCGUGUGAUUUUAAUAGCUUCUUAGGAGAAUG

Pair 2 (ara-UTP and dUTP extension assay):
Primer RNA: CAUUCUCCUAAGAAGCUAUUAAAAUCACA
Template RNA: UUUUUCCCAUGUGAUUUUAAUAGCUUCUUAGGAGAAUG

Pair 1 and 2 RNA oligos were synthesized by Integrated DNA Technologies. Primer RNAs were modified with a 5′ 6-FAM. RNA primers were annealed to templates at a ratio of 1:1.2. Annealing was performed in 2.5 mM HEPES pH 7.4, 2.5 mM potassium chloride, and 0.5 mM magnesium chloride by heating samples to 95 °C for 5 min, then gradually cooling to 25 °C for 75 min. Annealed RNAs were stored at -20°C.

Pair 3 (ara-CTP incorporation, pdb_00009BLF, EMD-44654):
Primer RNA: CAUUCUCCUAAGAAGCUAUUAAAAUCACA
Template RNA: AAAAAGGGUUGUGAUUUUAAUAGCUUCUUAGGAGAAUG

Pair 4: (ara-UTP and UTP incorporations, pdb_00009PYW, EMD-72038, pdb_00009PYZ, EMD-72053, pdb_00009PZ0, EMD-72054)
Primer RNA: CAUUCUCCUAAGAAGCUAUUAAAAUCACA
Template RNA: UUUUCCCAAUGUGAUUUUAAUAGCUUCUUAGGAGAAUG

Pair 3 and 4 RNA oligos were synthesized by Integrated DNA Technologies. Primer RNAs were modified with a 5′ 6-FAM. Primer and template RNAs were annealed at a 1:1 ratio in 2.5 mM HEPES pH 7.4, 2.5 mM potassium chloride, and 0.5 mM magnesium chloride by heating at 95 °C for 5 min, then gradually cooling to 25 °C for 75 min and RNA wase used immediately.

**SARS-CoV-2-catalyzed primer extensions with detection of $^{32}$P labeled products.**

To assemble elongation-competent SARS-CoV-2 RNA-dependent RNA polymerase (RdRp) complexes, nsp7, nsp8 and nsp12 subunits were prepared in enzyme dilution buffer containing 25 mM HEPES pH 7.5, 1 mM TCEP, and 20% glycerol. The volume of enzyme used in each reaction did not exceed one-tenth of the total reaction volume. SARS-CoV-2 polymerase complexes were freshly prepared before use where a mixture containing 0.5 µM nsp12, 1.5 µM nsp7, and 1.5 µM nsp8 was incubated with 1 µM double-stranded RNA (dsRNA) primer/template, 100 µM ATP, and 0.1 µCi/µL [α-$^{32}$P]-ATP for 5 min. at 30°C in 25 mM HEPES pH 7.5, 10 mM KCl, 2 mM $MgCl_2$, and 1 mM TCEP. Single nucleotide incorporation assays were initiated by adding either 0.1 µM CTP or UTP, or 1 mM of the corresponding nucleotide triphosphate analog (ara-CTP or ara-UTP) and allowed to proceed for 10 minutes before quenching with 50 mM EDTA. For chain termination experiments, reactions were performed in the presence of the next correct nucleotide substrate (1 µM UTP or 1 µM CTP), followed by quenching with 50 mM EDTA.

To evaluate inhibition of SARS-CoV-2 polymerase elongation by ara-CTP or ara-UTP, polymerase-RNA elongation complexes were assembled, and reactions were initiated with 0.1 µM CTP or UTP, along with increasing concentrations of ara-UTP or ara-CTP, respectively. These reactions were incubated for 10 minutes along with next correct nucleotide (1 µM UTP or 1 µM CTP) and then quenched with 50 mM EDTA.

Poliovirus polymerase reactions were carried out in a buffer containing 25 mM HEPES pH 7.5, 1 mM TCEP, 20% glycerol, and 1 µM poliovirus $3D^{pol}$ polymerase enzyme. The volume of enzyme added did not exceed one-tenth of the total reaction volume. Reactions were initiated by the addition of either 1 µM CTP or UTP, or 1 mM of the corresponding nucleotide analog (ara-CTP or ara-UTP), and allowed to proceed for 10 minutes before quenching with 50 mM EDTA. For chain termination assays, reactions were performed in the presence of the next correct nucleotide substrate (1 µM UTP or 1 µM CTP), followed by quenching with 50 mM EDTA.

For polyacrylamide gel electrophoresis (PAGE) analysis of reaction products, loading buffer contains 85% (v/v) formamide, 0.025% (w/v) bromophenol blue and 0.025% (w/v) xylene cyanol and an excess (50-fold) of unlabeled RNA oligonucleotide (trap strand) that is the exact same sequence to the $^{32}$P-labeled RNA oligonucleotide in the reaction (5'-CCGGGCGGCACU or 5'-CCGGGCGGCAUC for ara-CTP and ara-UTP). Having trap strand ensures complete separation and release of the $^{32}$P-lableled RNA oligonucleotide prior to gel electrophoresis. An equal volume of loading buffer was added to quenched reaction mixtures and heated to 90°C for 10 min prior to loading 5 µL of sample on a denaturing 20% polyacrylamide gel containing 1X TBE (89 mM Tris base, 89 mM boric acid, and 2 mM EDTA) and 7 M urea. Gels were visualized by using PhosphorImager (GE) and quantified by using ImageQuant TL software (GE). All gels shown are representative, single experiments that have been performed at least three to four times. In all cases, values for parameters measured during individual trials were within the limits of the error reported. Data were fit by either linear or nonlinear regression using the program GraphPad Prism v7.03 (GraphPad Software Inc.).

**High-throughput magnetic tweezers**

High-throughput magnetic tweezers apparatus.

The high-throughput magnetic tweezers used in this study have been described in detail elsewhere.[8] Shortly, a pair of vertically aligned permanent magnets (5 mm cubes, SuperMagnete, Switzerland) separated by a 1 mm gap are positioned above a flow chamber that is mounted on a custom-built inverted microscope. The vertical position and rotation of the magnets are controlled by two linear motors, M-126-PD1 and C-150 (Physik Instrumente PI, GmbH and Co. KG, Karlsruhe, Germany), respectively. The field of view is illuminated through the magnets gap by a collimated LED-light source and is imaged onto a large chip CMOS camera (Dalsa Falcon2 FA-80–12 M1H, Stemmer Imaging, Germany) using a 60× oil immersion objective (CFI Plan Apo $\lambda$, NA 1.4, Nikon, Germany) and an achromatic doublet tube lens of 200 mm focal length and 50 mm diameter (Qioptic, Germany). To control the temperature, we used a system previously described[6]. Shortly, a flexible resistive foil heater with an integrated 10 MΩ thermistor (HT10K, Thorlabs) is wrapped around the microscope objective and further insulated by several layers of Kapton tape (KAP22-075, Thorlabs). The heating foil is connected to a PID temperature controller (TC200 PID controller, Thorlabs) to adjust the temperature within ~0.1°C. The magnetic tweezers assay force calibration was described in detail in Ref. All data were acquired at 58 Hz camera frame rate.

Flow cell assembly.

The fabrication procedure for flow cells has been previously described[7]. To summarize, we sandwiched a double layer of Parafilm by two #1 coverslips, the top one having one hole at each end serving as inlet and outlet, the bottom one being coated with a 0.01% m/V nitrocellulose in amyl acetate solution. The flow cell is mounted into a custom-built holder and rinsed with ~1 ml of 1× phosphate-buffered saline (PBS). 3 µm diameter polystyrene reference beads are attached to the bottom coverslip surface by incubating 100 µl of a 1:1000 dilution in PBS of (LB30, Sigma Aldrich, stock concentration: $1.828*10^{11}$ particles per milliliter) for ~3 min. The tethering of the magnetic beads by the RNA hairpin construct relies on a digoxygenin/anti-digoxygenin and biotin-streptavidin attachment at the coverslip surface and the magnetic bead, respectively. Therefore, following a thorough rinsing of the flow cell with PBS, 50 µl of anti-digoxigenin (50 µg/ml in PBS) is incubated for 30 min. The flow cell was flushed with 1 ml of 10 mM Tris, 1 mM EDTA pH 8.0, 750 mM NaCl, 2 mM sodium azide buffer to remove excess of anti-digoxigenin followed by rinsing with another 0.5 ml of 1× TE buffer (10 mM Tris, 1 mM EDTA pH 8.0 supplemented with 150 mM NaCl, and 2 mM sodium azide). The surface is then passivated by incubating bovine serum albumin (BSA, New England Biolabs, 10 mg/ml in PBS and 50% glycerol) for 30 min and rinsed with 1× TE buffer.

Single-molecule RNA polymerase activity experiments.

For experiments with SARS-COV-2 polymerase, 20 µL of streptavidin-coated Dynal Dynabeads M-270 streptavidin-coated magnetic beads (Thermo Fisher Scientific) was mixed with ~0.1 ng of RNA hairpin (total volume 40 µL) and incubated for ~5 min before rinsing with ~2 mL of 1× TE buffer to remove any unbound RNA and the magnetic beads in excess. RNA tethers were sorted for functional hairpins by looking for the characteristic jump in extension of the correct length (~0.6 µm at 30 pN) due to the sudden opening of the hairpin during a force ramp experiment. The flow cell was subsequently rinsed with 0.5 mL reaction buffer (50 mM HEPES pH 7.9, 10 mM DTT, 2 µM EDTA, and 5 mM $MgCl_2$). After starting the data acquisition at a force that would keep the hairpin open, 100 µL of reaction buffer containing 0.6 µM of nsp12, 1.8 µM of nsp7 and nsp8, NTPs and ara-NTPs (if required) were flushed in the flow cell to start the reaction. The experiments were conducted at a constant force as indicated for a duration of 30 to 60 minutes. The camera frame rate was fixed at 58 Hz the temperature set to 25˚C. A custom written LabVIEW routine controlled the data acquisition and the (x-, y-, z-) positions analysis/tracking of both the magnetic and reference beads in real time. Mechanical drift correction was performed by subtracting the reference bead position to the magnetic bead position and by applying an autofocus.

For the experiments with poliovirus polymerase, 20 µL of streptavidin-coated Dynal Dynabeads M-270 streptavidin-coated magnetic beads (Thermo Fisher Scientific) was mixed with ~0.1 ng of dsRNA (total volume 40 µL) and incubated for ~5 min before rinsing with ~2 mL of 1× TE buffer to remove any unbound RNA and the magnetic beads in excess. RNA tethers were sorted for functional tethers by looking for the change in extension of the correct length (~0.8 - 1 µm at 30 pN) when the force was ramped from a lower to higher regime. The flow cell was subsequently rinsed with 0.5 mL reaction buffer (50 mM HEPES pH 7.9, 5 mM $MgCl_2$, 0.01% Triton X-100, 5% Superase RNase inhibitor (Life Technologies)). After starting data acquisition, the tethers were kept at a constant force of 30 pN, when 100 µL of reaction buffer containing 1.44 µM of poliovirus polymerase, the indicated concentration of NTPs and ara-NTP (if required) were flushed in the flow cell to start the reaction. The experiments were conducted for 60 to 75 mins with same software mentioned at a constant acquisition frequency of 58 Hz.

Magnetic tweezers data processing:

The replication activity of SARS-CoV-2 RNA polymerase converts the tether from ssRNA to dsRNA, which concomitantly decreases the end-to-end extension of the tether. The change in extension measured in micron was subsequently converted into replicated nucleotides $N_R$ using the following equation:

$$N_R(F) = N.\frac{L_{ss}(F)-L_{meas}(F)}{L_{ss}(F)-L_{ds}(F)} \qquad (1)$$

In the case of poliovirus, the polymerase unwinds the dsRNA to ssRNA which leads to increasing the length of the tether. To convert the position of the bead to the number of transcribed nucleotides $N_R$ , we use the linear-interpolation formula:

$$N_R(F) = N.\frac{L_{meas}(F) - L_{ds}(F)}{L_{ss}(F) - L_{ds}(F)} \quad (2)$$

where $L_{meas}(F)$, $L_{ss}(F)$ and $L_{ds}(F)$ are the measured extension during the experiment, the extension of an ssRNA and a dsRNA construct, respectively, experiencing a force F, and N the number of nucleotides of the ssRNA template. The traces were then filtered using a Kaiser-Bessel low-pass filter with a cutoff frequency at either 2 Hz or 0.5 Hz, when using either the open RNA hairpin or the dsRNA construct, respectively. As previously described[8], a dwell time analysis was performed by scanning the filtered traces with non-overlapping windows of 10 nt to measure the time (coined throughout the manuscript as dwell time) for SARS-CoV-2 or poliovirus polymerase to incorporate ten successive nucleotides. The dwell times of all the traces for a given experimental condition were assembled and further analyzed using a maximum likelihood estimation (MLE) fitting routine to extract the parameters from the stochastic-pausing model.

<u>SARS-CoV-2/poliovirus polymerase replication time and product length analysis.</u>

To extract the product length and replication of the replication complex, only the traces where the beginning and the end could clearly be distinguished and for which the tether did not rupture for five minutes following the last observed replication activity were considered. We represented the mean, as well as one standard deviation of the mean from 1000 bootstraps as error bars.

<u>Stochastic-pausing model.</u>

We use the kinetic model previously described[8,9]. The general form of the dwell time distribution without specifying how the pauses are connected to the nucleotide addition pathway is:

$$p_{dw}(t) \propto p_{na}\Gamma\left(t;\, N_{dw}, \frac{1}{k_{na}}\right) + Q(t)\left(\sum_{n=1}^{N_{sp}} p_n k_n e^{-k_n t}\right) \quad (3)$$

Without power law distribution, while when employing a power law distribution, we have:

$$p_{dw}(t) \propto p_{na}\Gamma\left(t;\, N_{dw}, \frac{1}{k_{na}}\right) + Q(t)\left(\sum_{n=1}^{N_{sp}} p_n k_n e^{-k_n t} + \frac{a_{bt}}{2(1+t/1s)^{3/2}}\right) \quad (4)$$

In the above expression, the gamma function in the first term contributes the portion $p_{na}$ of dwell times that originate in the polymerase crossing the dwell time window of size $N_{dw}$ base pairs without pausing; the second term is a sum of contributions originating in pause-dominated transitions, each contributing a fraction of $p_n$ dwell times. The backtracked asymptotic term needs to be regularized for times shorter than the diffusive backtrack step. We have introduced a regularization at 1 s, but the precise timescale does not matter, as long as it is set within the region where the exponential pauses dominate over the backtrack. From left to right, each term of equation 2 is dominating the distribution for successively longer dwell times.

A cut-off factor Q(t) for short times is introduced to account for the fact that the dwell time window includes $N_{dw}$ nucleotide-addition steps,

$$Q(t) = \frac{(tk_{na}/N_{dw})^{N_{dw}-1}}{1+(tk_{na}/N_{dw})^{N_{dw}-1}} \quad (5)$$

The dependence of the extracted fit parameters on these cut-offs is negligible, as long as they are introduced in regions where the corresponding term is sub-dominant. Here, the cut is placed under the center of the elongation peak, guaranteeing that it is placed where pausing is sub-dominant.

<u>Maximum likelihood estimation (MLE).</u>

The normalized version of Equation (3) is the dwell time distribution fit to the experimentally collected dwell-times $\{t_i\}_i$ by minimizing the likelihood function:

$$L = -\sum_i \ln p_{dw}(t_i) \quad (6)$$

with respect to rates and probabilistic weights.

Dominating in a dwell time window versus dominating in one step.

The fractions $p_n$ represent the probability that a particular rate $k_n$ dominates the dwell time. We want to relate this to the $P_n$ probability that a specific exit rate dominates within a 1-nt transcription window. Assuming we have labeled the pauses so that $k_{n-1} > k_n$, we can relate the probability of having rate $n$ dominating in $N_{dw}$ steps to the probability of having it dominate in one step through:

$$Lp_n = (\sum_{m=0}^{n} P_m)^{N_{dw}} - (\sum_{m=0}^{n-1} P_m)^{N_{dw}}, p_0 = p_{na} = P_{na}^{N_{dw}} = P_0^{N_{dw}} \quad (7)$$

The first term in equation (7) represents the probability of having no pauses longer than the $n^{th}$ pause in the dwell time window, and the second term represents the probability of having no pauses longer than the $(n-1)^{th}$ pause. The difference between the two terms is the probability that the $n^{th}$ pause will dominate. This can be inverted to yield a relation between the single-step probabilities ($P_n$) and the dwell time window probabilities ($p_n$):

$$P_n = (\sum_{m=0}^{n} p_m)^{1/N_{dw}} - (\sum_{m=0}^{n-1} p_m)^{1/N_{dw}}, P_0 = p_0^{1/N_{dw}} \quad (8)$$

This relationship has been used to relate our fits over a dwell time window to the single-step probabilities.

**Fluorescence-based polymerase activity gel assay for analog incorporation and extension**

1 µM nsp12, 2 µM nsp8, 2 µM nsp7 and 250 nM duplex RNA were used for primer extension reactions in 10 mM Tris-Cl pH 8.0, 100 mM potassium glutamate, 2 mM MgCl2 and 1 mM DTT. The proteins were mixed and incubated at 25 °C for 15 min, then RNA duplex was added and incubated at 25 °C for another 15 min. RNA pair 1 was used for CTP analog incorporations while pair 2 was used for UTP analog incorporations. Reactions were initiated by the addition of one of the NTP analogs at a final concentration of 40 µM. Analogs used include arabinofuranosylcytosine triphosphate (ara-CTP, Jena Biosciences), arabinofuranosyluridine triphosphate (ara-UTP, Jena Biosciences), deoxycytidine triphosphate (dCTP, New England Biolabs), deoxyuridine triphosphate (dUTP, New England Biolabs). The direct addition of all four natural NTPs served as a positive control for the reactions. Incorporation of nucleotide analogs proceeded for 3 min at 25 °C. Next, three NTPs other than the one that corresponds to the analog were added into the reaction at 25 °C (each nucleotide, 40 µM final concentration). Reaction timepoints were taken at 20 s, 1 min, 3 min, 7 min, 15 min, 30 min, and 60 min. Timepoint samples were quenched by adding two volumes of sample loading buffer (95% (v/v) formamide, 4 mM ethylenediaminetetraacetic acid (EDTA) and 0.75 mM bromophenol blue). Each reaction was heated at 95°C before loading on a TBE-urea-PAGE gel (15 % polyacrylamide, 8 M urea). Gels were imaged on Typhoon FLA 9000 imager (GE Healthcare) to detect 6-FAM fluorescence.

**Single-particle cryo-electron microscopy for structure determination**

SARS-CoV-2 polymerase araCMP-araCTP structure (pdb_00009BLF, EMD-44654). Proteins were assembled in 25 mM HEPES pH 7.5, 100 mM sodium chloride, 2 mM magnesium chloride, and 2 mM DTT at room temperature for 15 min. RNA duplexes were then added to assembled proteins and incubated for another 15 min. The complex were then concentrated to 40 µL. Immediately before freezing the sample, 2 µL of 800 µM ATP and araCTP were added into the complex. Samples were prepared at final concentration of 4 mg/mL protein with a protein-RNA ratio of 2:2:1:1.2 nsp7:nsp8:nsp12:duplex RNA.

Samples were loaded on UltraAuFoil R1.2/1.3 300 mesh grids (Quantifoil) after having been glow discharged for 20 s with 20 mA in air using a GloQube Plus (Quorom). Before freezing, 6 mM 3-([3-cholamidopropyl] dimethylammonio)-2-hydroxy-1-propanesulfonate (CHAPSO, final concentration) was added into the sample. 3 µL of sample was spotted onto grids for double-sided blotting in a Vitrobot Mark IV (ThermoFisher Scientific) at 4 °C and 100% humidity with -14 blot force and 10 s blot time before plunge freezing in liquid ethane.

Data were collected on a Talos Arctica 200 kV transmission electron microscope (ThermoFisher Scientific). Movies were collected with no tilt at 79,000x magnification and 1.064 Å pixel size, with a GIF quantum energy filter set to a 20 eV slit width and a K3 direct electron detector (Gatan) in CDS mode. The focus ranged from −0.5 to −2.0 µm. The total dose of each movie was 60 e⁻/Å².

Data were processed using CryoSPARC[10]. Movies were motion corrected using patch motion correction. After CTF estimation and particle picking, 1,306,642 particles were extracted with a box size of 256 pixels. The particles were subjected to one round of 2D classification. Three ab-initio models were generated.

Heterogeneous refinement was performed to re-classify particles and one of the maps was refined using non-uniform refinement. This particle stack was used to generate two new ab-initio reconstructions, one of which was subjected to 3D variability analysis and heterogeneous refinement resulting in a final map from 158,606 particles. A coordinate model was built and validated using PHENIX[11], COOT[12], and ISOLDE[13] and using 7UOE.pdb[14] as a starting mode.

SARS-CoV-2 polymerase araUMP (pdb_00009PYW, EMD-72038), araUMP-UTP (pdb_00009PYZ, EMD-72053), and UMP-UMP (pdb_00009PZ0, EMD-72054) structures

Proteins were assembled in 10 mM Tris-Cl pH 8, 100 mM potassium glutamate, 2 mM magnesium chloride, and 1 mM DTT and assembly was allowed to proceed for 15 min at room temperature. Duplex RNA was then added to the complex and allowed to assemble for an additional 15 min at room temperature. The complexes were then concentrated to 40 µL. For ara-UTP incoporations, 16 µL 400 µM ara-UTP was added to the complexes and incubated for 10 min. Alternatively, 16 µL of 400 µM UTP was added and incubated for 10 min. All samples were concentrated to 40 µL after nucleotide incorporations. All three samples contained a final concentration of 8 mg/ml protein with a protein-RNA ratio of 2:2:1:1.2 nsp7:nsp8:nsp12:duplex RNA.

UltraAuFoil R1.2/1.3 300 mesh grids (Quantifoil) were glow discharged in air in a GloQube Plus (Quorom) for 30 s and 20 mA. 6 mM CHAPSO detergent (final concentration) was added into the sample. For the araUMP-UTP dataset (pdb_00009PYZ, EMD-72053), 1 µL of 800 µM UTP was added immediately before adding CHAPSO and spotting onto grids. 3 µL of sample was added on the grids for double-sided blotting in a Vitrobot Mark IV (ThermoFisher Scientific at 4 °C and 100% humidity. The samples blotted with -7 blot force, 4 s blot time and plunge freezing in liquid ethane. All grids were screened on a Talos Arctica 200 kV transmission electron microscope (ThermoFisher Scientific) with no tilt by using a GIF quantum energy filter with a 20 eV slit width and K3 direct electron detector (Gatan) in CDS mode. Data were collected at 79,000x magnification, 1.064 Å pixel size, and 55 $e^{-}/Å^2$ total dose per movie. The focus ranged from −0.5 to −2.0 µm.

Data were processed in CryoSPARC[10] using patch motion correction, CTF estimation and particle picking to extract 1,778,717 initial particles for the ara-UMP structure, 1,255,633 initial particles for the ara-UMP-UTP structure, and 1,928,958 initial particles for the UMP-UMP structure each with a box size of 256. The particles went through two rounds of 2D classification for the ara-UMP structure and one round of 2D classification in araUMP-UTP and UMP-UMP structures. After ab-initio reconstruction, heterogeneous refinement, and non-uniform refinement, 621,798 particles for the araUMP structure, 775,227 particles for the araUMP-UTP structure, and 740,997 particles for the UMP-UMP structure were used to reconstruct final maps. The models were built and validated using PHENIX[11], COOT[12], and ISOLDE[13] using the ara-CMP-ara-CTP (pdb_00009BLF, EMD-44654) as a starting model.

**Poliovirus 3D$^{pol}$ polymerase stopped-flow experiments**

Pre-steady state nucleotide incorporation stopped-flow experiments were performed as described previously.[1,2] All reactions were performed at 30°C using a Model SF-300X stopped-flow apparatus (Kintek Corp., Austin, TX) equipped with a water-bath. The poliovirus 3D$^{Pol}$ polymerase was diluted into enzyme buffer (50 mM HEPES, pH 7.5, 1 mM TCEP, and 20% glycerol) immediately prior to use. One-twentieth volume of enzyme was added to each reaction. Reactions were performed by incubating 1 µM enzyme with 1 µM sym/sub RNA primer-template (0.5 µM duplex) in 25 mM HEPES pH 7.5, 1 mM TCEP, 60 µM zinc chloride, and 5 mM magnesium chloride at room temperature for 3 minutes. For reactions with a final NTP concentration higher than 1 mM, the amount of free $Mg^{+2}$ was kept constant by increasing the amount of magnesium chloride in the reaction to 5 mM plus the NTP concentration above 1 mM. Reactions were allowed to equilibrate to 30°C and then rapidly mixed with the indicated NTP or ara-NTP.

The sym/sub RNA primer-template substrates have as the first templating base (0 position) either a G when using either CTP and ara-CTP or an A when using either UTP and ara-UTP. There is also a fluorescent reporter nucleotide at the +1 templating position, either 2-aminopurine (2AP) or pyrrolo-C (PC) for CTP and UTP incorporation respectively. For 2AP, the excitation wavelength was 313 nm and fluorescence emission was monitored by using a 370 nm cut-on filter (model E370LP, Chroma Technology Corp., Rockingham, VT). For PC, the excitation wavelength was 350 nm and fluorescence emission was monitored by using a 450 nm cut-on filter (model HW440LP, Chroma Technology Corp., Rockingham, VT). Single CTP or ara-CTP incorporation into the sym/sub-G RNA primer-template results in a decrease in 2AP fluorescence. Conversely, single UTP or ara-UTP incorporation into the sym/sub-pyrroloC RNA primer-template results in an increase in PC fluorescence.

To determine the apparent dissociation constant ($K_{d,app}$) and polymerization rate constant ($k_{pol}$) for ara-UTP and ara-CTP, a range of substrate concentrations was used. Specifically, ara-UTP was tested at concentrations from 25 to 1250 µM, and ara-CTP from 10 to 100 µM. For comparison, the corresponding natural nucleotides were evaluated over similar ranges: UTP from 75 to 1500 µM and CTP from 7.5 to 90 µM. For each experiment, at least six fluorescence traces were averaged. Relative fluorescence was plotted as a function of time and fit to a single-exponential equation (eq. 9).

$$F = Ae^{-kobs*t} + C \qquad (9)$$

where $A$ is the amplitude, $k_{obs}$ is the observed rate constant, and $C$ is the end point. Values for $k_{pol}$, the maximal rate constant for single-nucleotide incorporation, and $K_{d,app}$, the apparent dissociation constant for NTP, were obtained by first determining $k_{obs}$ by using equation (9) for a range of NTP concentrations and then fitting plots of $k_{obs}$ dependence on NTP concentration to a hyperbolic equation (eq. 10).

$$kobs = \frac{kpol*[NTP]}{Kd,app+[NTP]} \qquad (10)$$

Kinetic data were fit by nonlinear regression using the program GraphPad Prism 10 (GraphPad Software, Inc.).

### SUPPLEMENTAL TABLES

| NTP conc. (μM) | 500 A/C/G/UTP | 500 A/C/G/UTP | 500 A/G/UTP, 50 CTP |
|---|---|---|---|
| ara-CTP conc. (μM) | 0 | 500 | 500 |
| Number of traces | 161 | 124 | 154 |
| Product length ± std (nt) | 1,039 ± 8 | 1,052 ± 12 | 1,065 ± 15 |
| Total replication time ± std (s) | 22.6 ± 0.95 | 26.9 ± 0.78 | 150.6 ± 17 |
| Dwell time distribution | | | |
| Number of dwell times | 14,840 | 10,485 | 15,095 |
| Nucleotide addition rate ± std (1/s) | 75.8 ± 0.7 | 79.02 ± 0.6 | 67.7 ± 0.6 |
| Pause1 exit rate ± std (1/s) | 4.99 ± 0.17 | 4.19 ± 0.1 | 3.72 ± 0.06 |
| Pause2 exit rate ± std (1/s) | 0.99 ± 0.18 | 0.40 ± 0.10 | 0.74 ± 0.07 |
| Ara-CTP pause exit rate ± std (1/s) | NA | 0.003 ± 0.002 | 0.005 ± 0.0005 |
| Pause1 probability ± std | 0.05 ± 0.002 | 0.05 ± 0.001 | 0.09 ± 0.002 |
| Pause2 probability ± std | 0.001 ± 0.0001 | 0.0020 ± 0.0001 | 0.0020 ± 0.0002 |
| Ara-CTP pause probability ± std | NA | 0.0003 ± 0.00009 | 0.001 ± 0.0001 |
| Backtrack probability ± std | 0.0008 ± 0.0001 | NA | NA |
| Figures | Fig. 2, S2 | Fig. S2 | Fig. 2 |

**Table S1: Extension of a ssRNA template by the SARS-CoV-2 RNA polymerase.** The table lists data collection statistics and refined model parameters for polymerase RNA extensions on single-stranded templates with natural NTPs or with the inclusion of ara-CTP. All experiments were run at 25˚C and 25 pN of force. NA, not applicable.

| | | | | |
|---|---|---|---|---|
| EMDB | 44654 | 72038 | 72053 | 72054 |
| PDB | 9BLF | 9PYW | 9PYZ | 9PZ0 |
| **Data collection** | | | | |
| Voltage (kV) | 200 | 200 | 200 | 200 |
| Dose Rate ($e^-$/pixel/sec) | 14.2 | 14.0 | 13.8 | 14.0 |
| Exposure Time (sec) | 4.82 | 4.45 | 4.52 | 4.45 |
| Electron Exposure ($e^-/Å^2$) | 60 | 55 | 55 | 55 |
| Frames (no.) | 60 | 55 | 55 | 55 |
| Defocus Values (µm) | -0.5 - -2.0 | -0.5 - -2.0 | -0.5 - -2.0 | -0.5 - -2.0 |
| Nominal Magnification | 79,000 | 79,000 | 79,000 | 79,000 |
| Pixel Size (Å) | 1.064 | 1.064 | 1.064 | 1.064 |
| Symmetry Imposed | C1 | C1 | C1 | C1 |
| Movies Collected (no.) | 4,230 | 3,479 | 4,501 | 4,999 |
| Initial Particle Images (no.) | 1,306,642 | 1,778,717 | 1,255,633 | 1,928,958 |
| Final Particle Images (no.) | 158,606 | 621,798 | 775,227 | 740,997 |
| Map Resolution (Å) | 3.3 | 3.1 | 3.1 | 3.0 |
| **Coordinate model refinement** | | | | |
| Initial Models Used (PDB) | 7UOE | 9BLF | 9BLF | 9BLF |
| Non-hydrogen Atoms | 12,364 | 12,118 | 12,148 | 12,032 |
| Protein Residues | 1,375 | 1,369 | 1,369 | 1,366 |
| Nucleic Acid Residues | 64 | 57 | 57 | 55 |
| Other Atoms | 3 $Mg^{2+}$, 2 $Zn^{2+}$ | 1 $Mg^{2+}$, 2 $Zn^{2+}$ | 2 $Mg^{2+}$, 2 $Zn^{2+}$ | 2 $Zn^{2+}$ |
| Ligands | 1 ara-CMP, 2 ara-CTP | 1 ara-UMP | 1 ara-UMP, 1 UTP | - |
| R.M.S. Deviations | | | | |
| Bond Lengths (Å) | 0.004 | 0.012 | 0.012 | 0.012 |
| Bond angles (°) | 0.942 | 1.889 | 1.902 | 1.911 |
| MolProbity[15] Score | 1.14 | 0.92 | 0.93 | 0.81 |
| Clashscore | 3.49 | 1.71 | 1.75 | 1.07 |
| Ramachandran Plot | | | | |
| Favored (%) | 98.46 | 98.38 | 98.09 | 98.23 |
| Allowed (%) | 1.54 | 1.62 | 1.91 | 1.77 |
| Outliers (%) | 0.00 | 0.00 | 0.00 | 0.00 |

**Table S2: cryo-EM data collection and model refinement parameters.** Map and model statistics for each of the four structures presented are tabulated. All data was collected on a Talos Arctica operating at 200 kV, 79,000x nominal magnification with a K3 direct electron detector (Gatan) in CDS counting mode. Map resolutions were determined from gold-standard Fourier shell correlations of two half maps at an correlation value of 0.143. Coordinate model quality was determined in PHENIX[11]. From left to right, the columns represent the structures of the SARS-CoV-2 polymerase complex bound to RNA with incorporated ara-CMP and incoming ara-CTP, ara-UMP, ara-UMP with an incoming UTP, or two incorporated UMP.

| Substrate | NTP | $K_{d,app}$(μM) | $k_{pol}$(s$^{-1}$) | $k_{pol}$ / $K_{d,app}$ (μM$^{-1}$ s$^{-1}$) | $k_{pol}$ / $K_{d,app}$ NTP/ara-NTP |
|---|---|---|---|---|---|
| sym/sub-G<br>5′ UCA**G**CCCGGG 3′<br>3′ GGGCCC**G**ACU 5′ | CTP | 22 | 110 | 5.0 | 50 |
| | ara-CTP[a] | 35 | 3.5 | 0.1 | |
| | ara-CTP[b] | 16 | 0.6 | 0.04 | |
| sym/sub-pyrolloC<br>5′ GC**A**UGGGCCCA 3′<br>3′ ACCCGGGU**A**CG 5′ | UTP | 150 | 140 | 0.90 | 90 |
| | ara-UTP | 230 | 1.9 | 0.01 | |

**Table S3: Kinetic parameters for 3D$^{pol}$ catalyzed nucleotide incorporation.** Templating base for single nucleotide incorporations are in bold. Fluorescent nucleotides 2AP and pyrrolo-C are colored blue in the sym/sub-G and sym/sub-pyrolloC RNA sequences. [a,b]Double exponential fits account for amplitude contributions from both fast and slow phases.

| NTP | Conc. (μM) | $k_{obs1}$ | $Amp_1$ | $k_{obs2}$ | $Amp_2$ |
|---|---|---|---|---|---|
| CTP | 7.5 | 25.79 | 1.2 | | |
| | 15 | 46.40 | 1.1 | | |
| | 30 | 64.85 | 1.1 | | |
| | 45 | 79.00 | 1.0 | | |
| | 60 | 78.00 | 1.6 | | |
| | 75 | 87.36 | 1.4 | | |
| | 90 | 90.17 | 1.2 | | |
| ara-CTP | 20 | 1.3 | 1.1 | 0.33 | 0.63 |
| | 40 | 1.8 | 1.3 | 0.42 | 0.68 |
| | 60 | 2.2 | 1.3 | 0.46 | 0.61 |
| | 80 | 2.4 | 1.3 | 0.50 | 0.57 |
| | 100 | 2.6 | 1.3 | 0.50 | 0.54 |
| UTP | 75 | 46.89 | -0.91 | | |
| | 150 | 70.15 | -0.94 | | |
| | 300 | 91.27 | -0.75 | | |
| | 450 | 103.47 | -1.24 | | |
| | 600 | 110.34 | -0.90 | | |
| | 750 | 113.56 | -1.42 | | |
| | 900 | 116.9 | -0.86 | | |
| | 1200 | 122.83 | -0.77 | | |
| | 1500 | 123.68 | -0.92 | | |
| ara-UTP | 25 | 0.16 | -1.05 | | |
| | 50 | 0.25 | -0.92 | | |
| | 100 | 0.57 | -0.96 | | |
| | 200 | 0.85 | -0.91 | | |
| | 500 | 1.38 | -0.92 | | |
| | 750 | 1.40 | -0.93 | | |
| | 1000 | 1.50 | -0.96 | | |
| | 1250 | 1.52 | -0.98 | | |

**Table S4: Raw kinetic values from stopped flow data.** Shown are the values obtained from fitting the stopped-flow fluorescence data to either a single or double exponential as shown in Fig. S12.

| NTP conc. (μM) | 500 A/C/G/UTP | 500 A/C/G/UTP | 500 A/C/G/UTP | 500 A/C/G/UTP |
|---|---|---|---|---|
| Ara-CTP conc. (μM) | 0 | 50 | 100 | 500 |
| Number of traces | 73 | 22 | 87 | 131 |
| Product length ± std (nt) | 3,014 ± 25 | 2,820 ± 31 | 2,796 ± 16 | 2,970 ± 51 |
| Total replication time ± std (s) | 243.7 ± 53.31 | 547.9 ± 308.3 | 707.1 ± 126.05 | 1,077.8 ± 138.05 |
| Dwell time distribution | | | | |
| Number of dwell times | 17,326 | 4,201 | 18,933 | 22,411 |
| Nucleotide addition rate ± std (1/s) | 24.1 ± 0.1 | 23.7 ± 0.2 | 23.1 ± 0.1 | 22.9 ± 0.1 |
| Pause1 exit rate ± std (1/s) | 2.02 ± 0.11 | 1.59 ± 0.17 | 1.89 ± 0.06 | 1.37 ± 0.04 |
| Pause2 exit rate ± std (1/s) | 0.45 ± 0.035 | 0.18 ± 0.005 | 0.18 ± 0.018 | 0.14 ± 0.02 |
| Ara-CTP pause exit rate ± std (1/s) | NA | 0.002 ± 0.0002 | 0.0016 ± 0.0003 | 0.0017 ± 0.0002 |
| Pause1 probability ± std | 0.018 ± 0.001 | 0.018 ± 0.002 | 0.018 ± 0.0008 | 0.023 ± 0.0006 |
| Pause2 probability ± std | 0.001 ± 0.001 | 0.00094 ± 0.0002 | 0.00069 ± 0.00007 | 0.0008 ± 0.0001 |
| Ara-CTP pause probability ± std | NA | 0.00022 ± 0.00005 | 0.00026 ± 0.00002 | 0.0008 ± 0.00004 |
| Backtrack probability ± std | 0.0004 ± 0.00006 | NA | NA | NA |

**Table S5. Extension of a dsRNA template by the poliovirus polymerase.** The table lists data collection statistics and refined model parameters for polymerase RNA extensions on double-stranded templates with natural NTPs or with the inclusion of ara-CTP. This table corresponds data presented in Fig. S3. All experiments were run at 25˚C and 30 pN of force. NA, not applicable. Elongation traces and dwell time log histograms are presented in Fig. S13.

| NTP conc. (µM) | 500 A/C/G/UTP | 500 A/C/G/UTP | 500 A/C/G/UTP | 500 A/C/G/UTP |
|---|---|---|---|---|
| Ara-UTP conc. (µM) | 0 | 50 | 100 | 500 |
| Number of traces | 73 | 97 | 97 | 87 |
| Product length (mean± std) nt | 3,014 ± 25 | 3,016 ± 17 | 3,033 ± 20 | 2,943 ± 63 |
| Total replication time (mean± std) s | 243.7 ± 53.31 | 315.8 ± 56.14 | 375.5 ± 66.15 | 1,182.2 ± 227.81 |
| Dwell time distribution | | | | |
| Number of dwell times | 17,326 | 24158 | 23095 | 11289 |
| Nucleotide addition rate ± std (1/s) | 24.1 ± 0.1 | 26.6 ± 0.1 | 26.8 ± 0.1 | 26.8 ± 0.2 |
| Pause1 exit rate ± std (1/s) | 2.02 ± 0.11 | 1.87 ± 0.11 | 1.52 ± 0.06 | 2.10 ± 0.10 |
| Pause2 exit rate ± std (1/s) | 0.45 ± 0.035 | 0.37 ± 0.033 | 0.09 ± 0.018 | 0.40 ± 0.06 |
| Ara-UTP pause exit rate ± std (1/s) | NA | 0.0035 ± 0.0013 | 0.0012 ± 0.00036 | 0.003 ± 0.0005 |
| Pause1 probability ± std | 0.018 ± 0.001 | 0.014 ± 0.0009 | 0.012 ± 0.0005 | 0.026 ± 0.0014 |
| Pause2 probability ± std | 0.001 ± 0.001 | 0.0018 ± 0.0001 | 0.0005 ± 0.00005 | 0.0018 ± 0.0001 |
| Ara-UTP pause probability ± std | NA | 0.00024 ± 0.00003 | 0.00017 ± 0.00003 | 0.00083 ± 0.00006 |
| Backtrack probability ± std | 0.0004 ± 0.00006 | NA | NA | NA |

**Table S6. Extension of a dsRNA template by the poliovirus polymerase.** The table lists data collection statistics and refined model parameters for polymerase RNA extensions on double-stranded templates with natural NTPs or with the inclusion of ara-UTP. This table corresponds data presented in Fig. S4. All experiments were run at 25˚C and 30 pN of force. NA, not applicable. Elongation traces and dwell time log histograms are presented in Fig. S14.

### SUPPLEMENTAL FIGURES

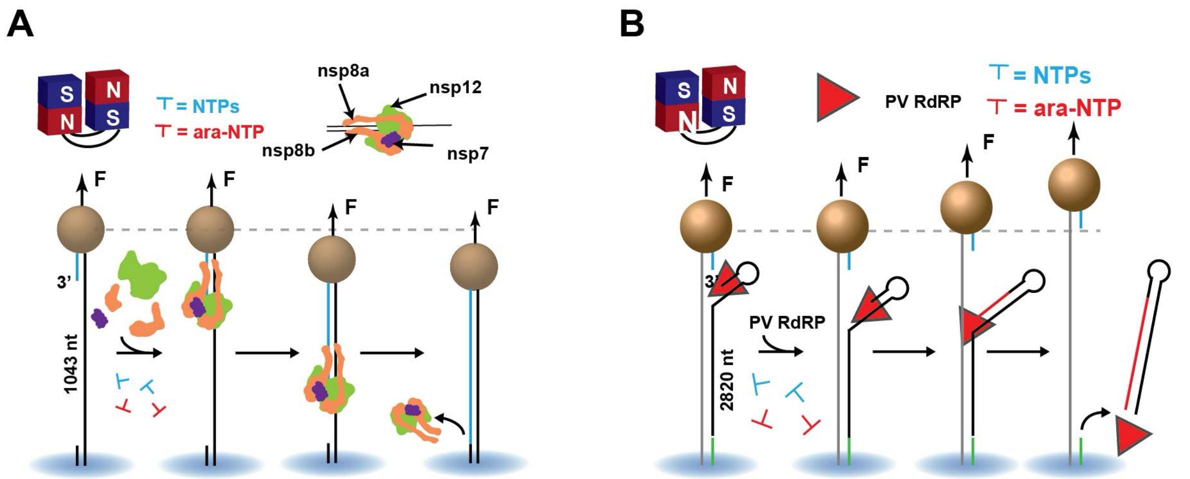


**Figure S1: Magnetic tweezer assay schematics. A)** Schematic of the magnetic tweezers assay to monitor SARS-CoV-2 polymerase RNA synthesis activity. **B)** Schematic of the magnetic tweezers assay to monitor poliovirus RNA polymerase RNA synthesis activity.

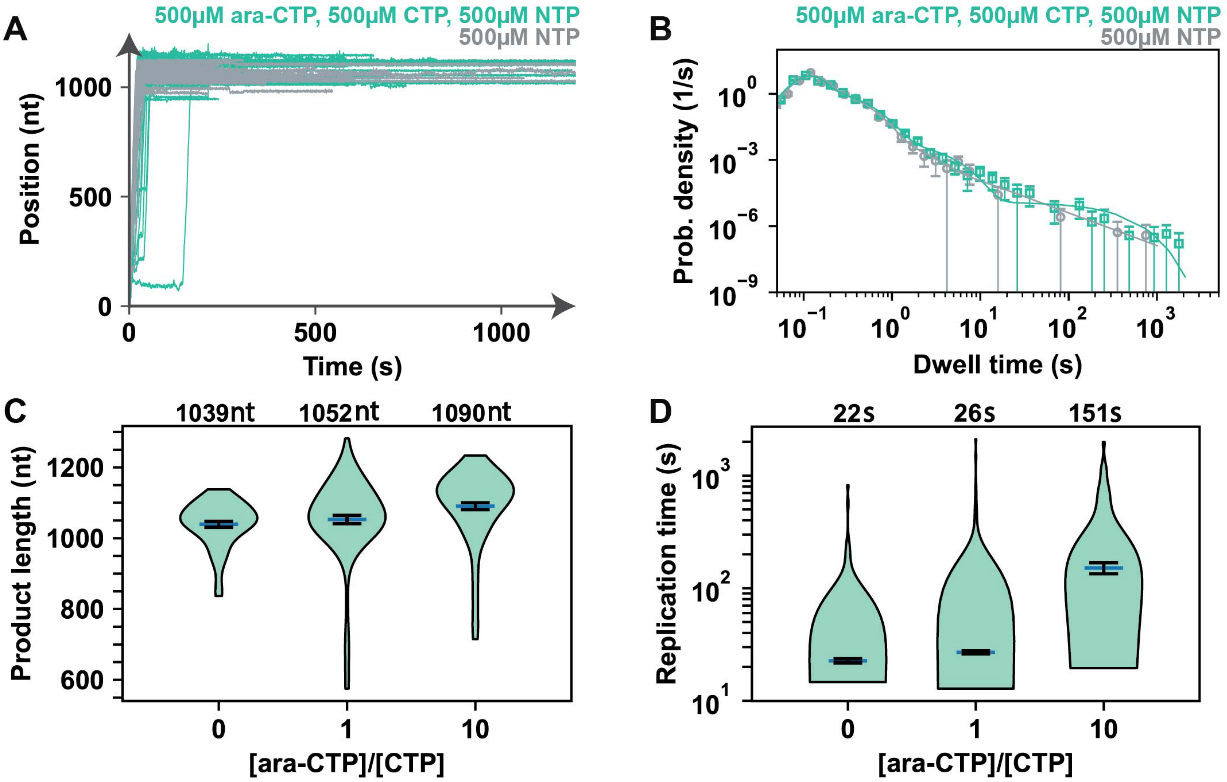


**Figure S2: ara-CTP induces pauses of long duration upon incorporation by the SARS-CoV-2 polymerase. A)** SARS-CoV-2 polymerase activity traces for either 0 μM ara-CTP and 500 μM NTP (grey), or 500 μM ara-CTP and 500 μM NTP (light green). **B)** Dwell time distributions of SARS-CoV-2 polymerase activity traces under varied reaction conditions, as illustrated in (A) with the same color shades. Fitting with ara-CTP pause was done for the conditions where ara-CTP was present in the reaction mixture. **C)** Product length of the SARS-CoV-2 polymerase over 1,043 nt long template as a function of the [ara-CTP]/[CTP] ratio. **D)** SARS-CoV-2 polymerase replication time as a function of the [ara-CTP]/[CTP] ratio. The mean values of the product length and replication time are indicated over the violin plot as a dark blue solid line flanked by two horizontal black lines representing one standard deviation error bars extracted from 1,000 bootstraps.

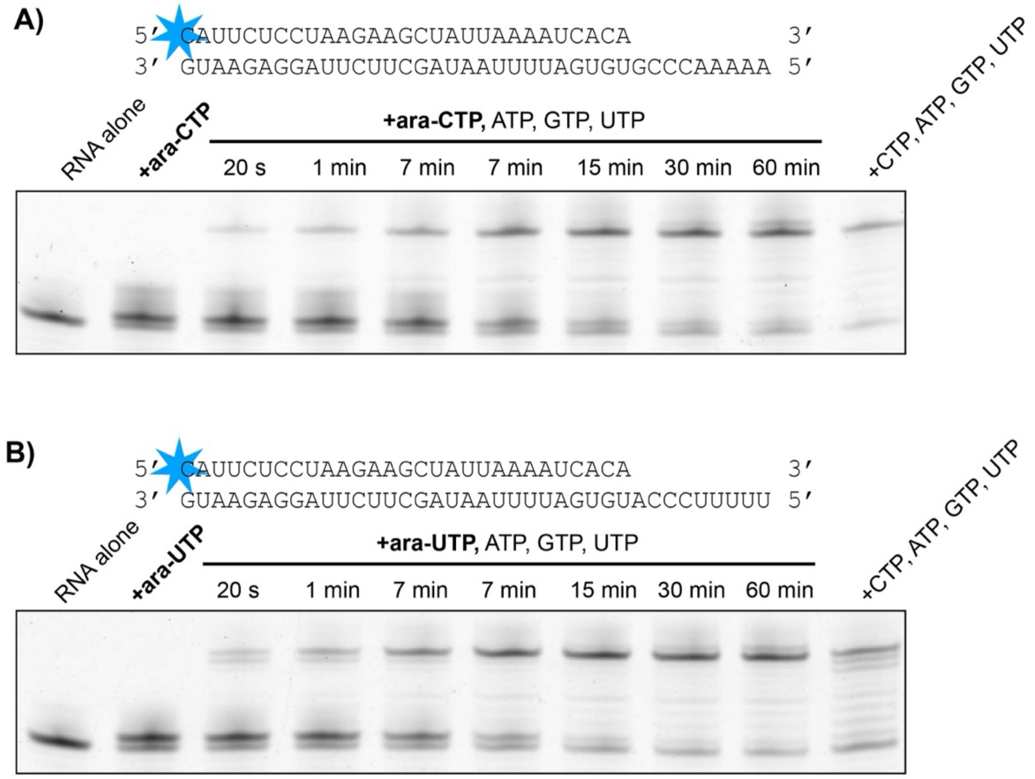


**Figure S3: ara-NTP Incorporation and chase assay.** Indicated primer template pairs were used to incorporate **A)** ara-CTP or **B)** ara-UTP and then the reactions were chased with the remaining nucleotides and followed over time. Primers are labeled with a 5’ 6-FAM tag and reactions are analyzed using denaturing urea-PAGE with FAM detection.

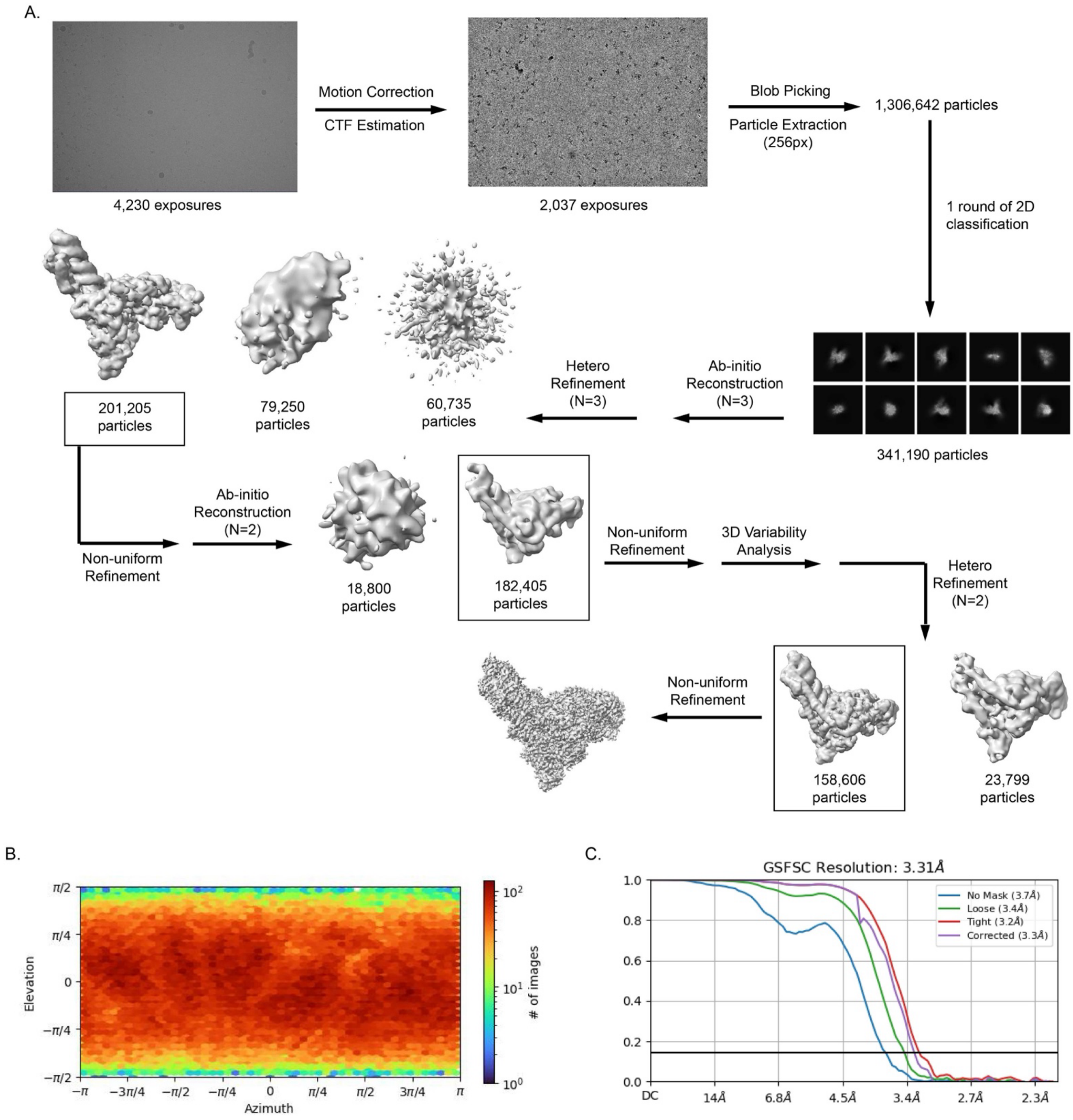


**Figure S4: Cryo-EM workflow and validation for SARS-CoV-2 RTC with incorporated ara-CMP and incoming ara-CTP. A)** Cryo-EM data were processed using cryoSPARC[10]. **B)** Orientation distribution plot for the final reconstructed map. **C)** Gold-standard Fourier shell correlation (GFFSC) of two independent half maps. The reported resolution is the frequence at a correlation of 0.143. Resulting reconstructed maps and coordinate models have been deposited as pdb_00009BLF, EMD-44654.

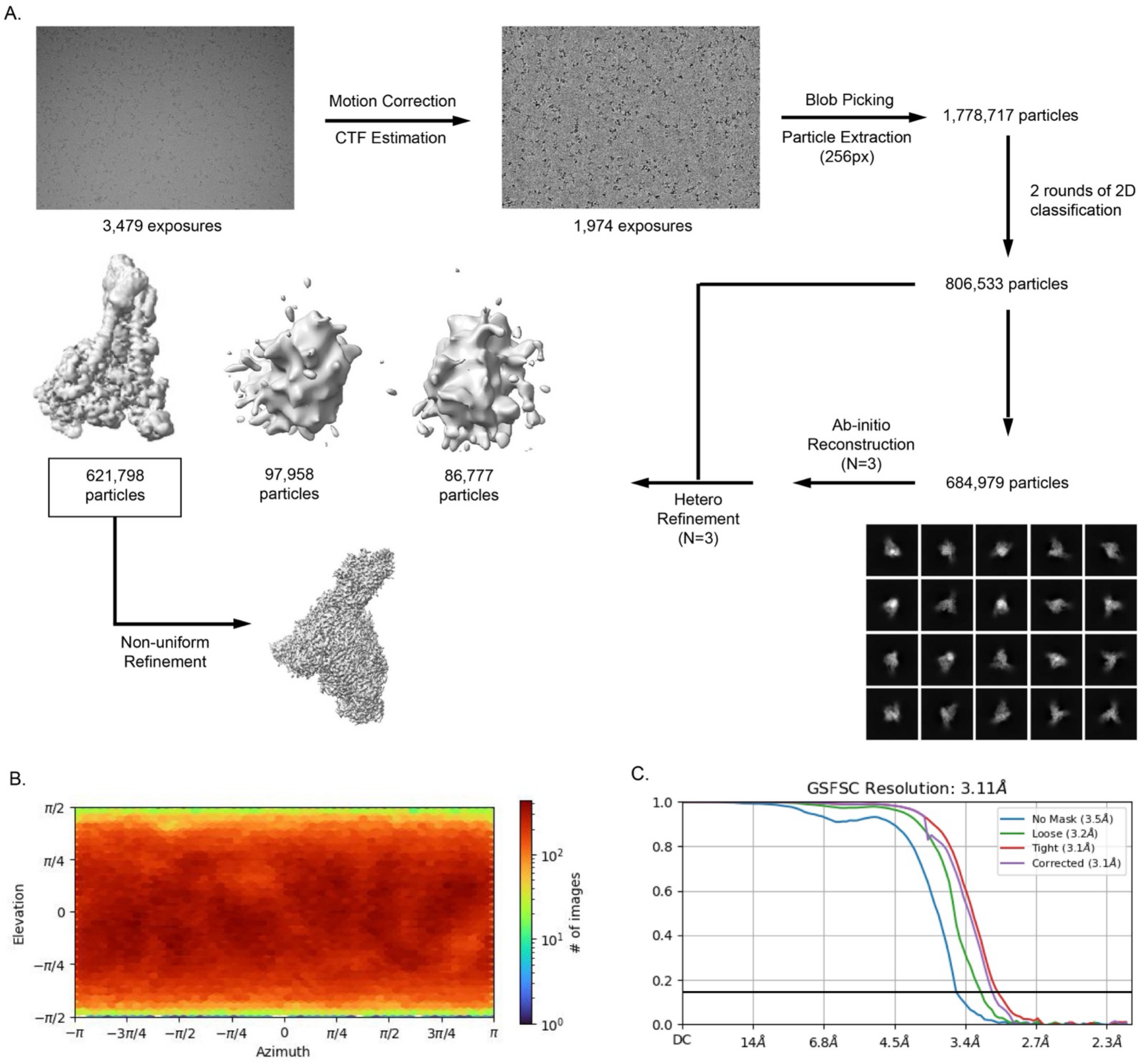


**Figure S5: Cryo-EM workflow and validation for SARS-CoV-2 RTC with incorporated ara-UMP.** Cryo-EM data were processed using cryoSPARC[10]. **B)** Orientation distribution plot for the final reconstructed map. **C)** Gold-standard Fourier shell correlation (GFFSC) of two independent half maps. The reported resolution is the frequence at a correlation of 0.143. Resulting reconstructed maps and coordinate models have been deposited as pdb_00009PYW, EMD-72038.

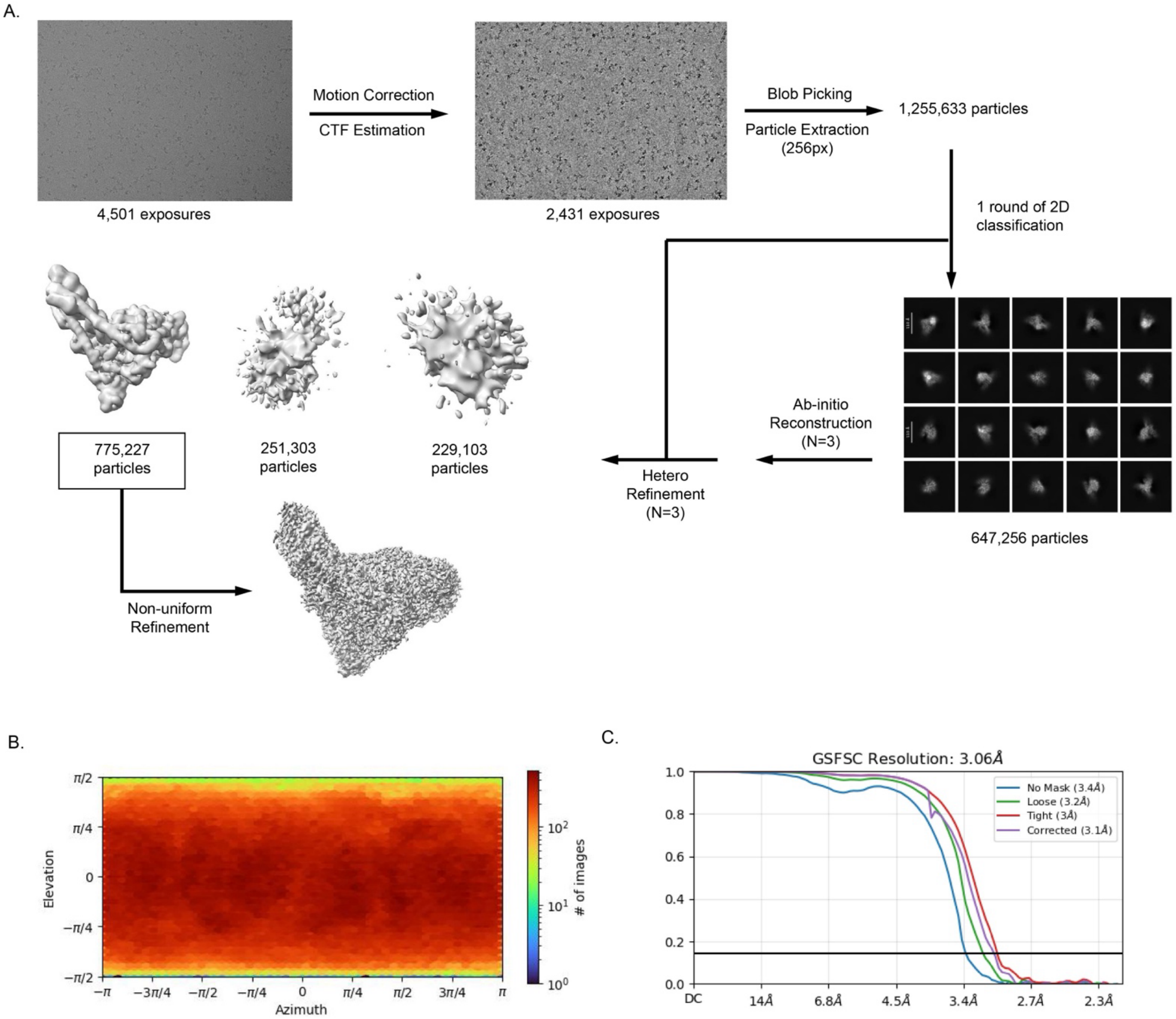


**Figure S6: Cryo-EM workflow and validation for SARS-CoV-2 RTC with incorporated ara-UMP and incoming UTP.** Cryo-EM data were processed using cryoSPARC[10]. **B)** Orientation distribution plot for the final reconstructed map. **C)** Gold-standard Fourier shell correlation (GFFSC) of two independent half maps. The reported resolution is the frequence at a correlation of 0.143. Resulting reconstructed maps and coordinate models have been deposited as pdb_00009PYZ, EMD-72053.

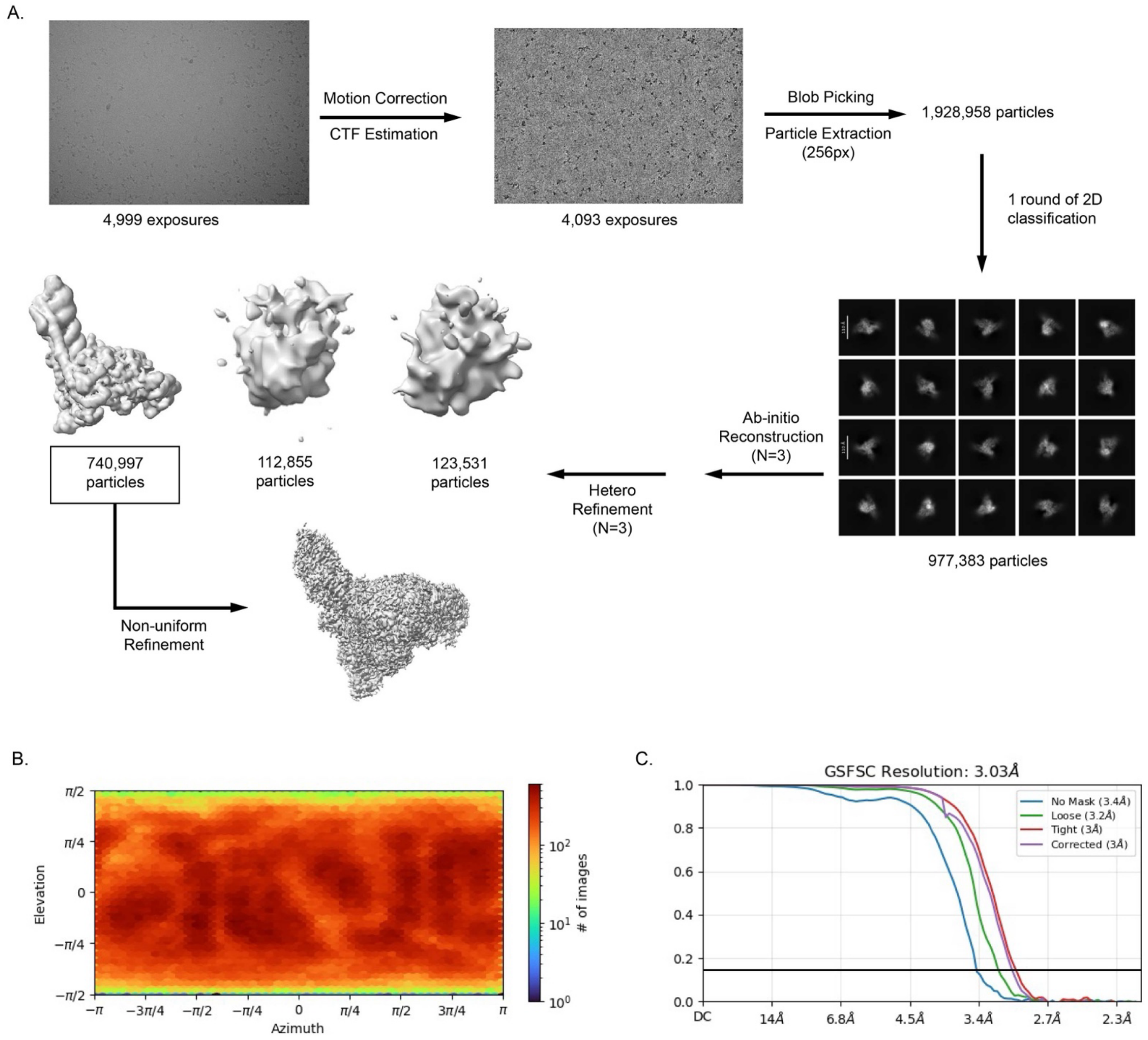


**Figure S7: Cryo-EM workflow and validation for SARS-CoV-2 RTC with incorporated UMP.** Cryo-EM data were processed using cryoSPARC[10]. **B)** Orientation distribution plot for the final reconstructed map. **C)** Gold-standard Fourier shell correlation (GFFSC) of two independent half maps. The reported resolution is the frequence at a correlation of 0.143. Resulting reconstructed maps and coordinate models have been deposited as pdb_00009PZ0, EMD-72054.

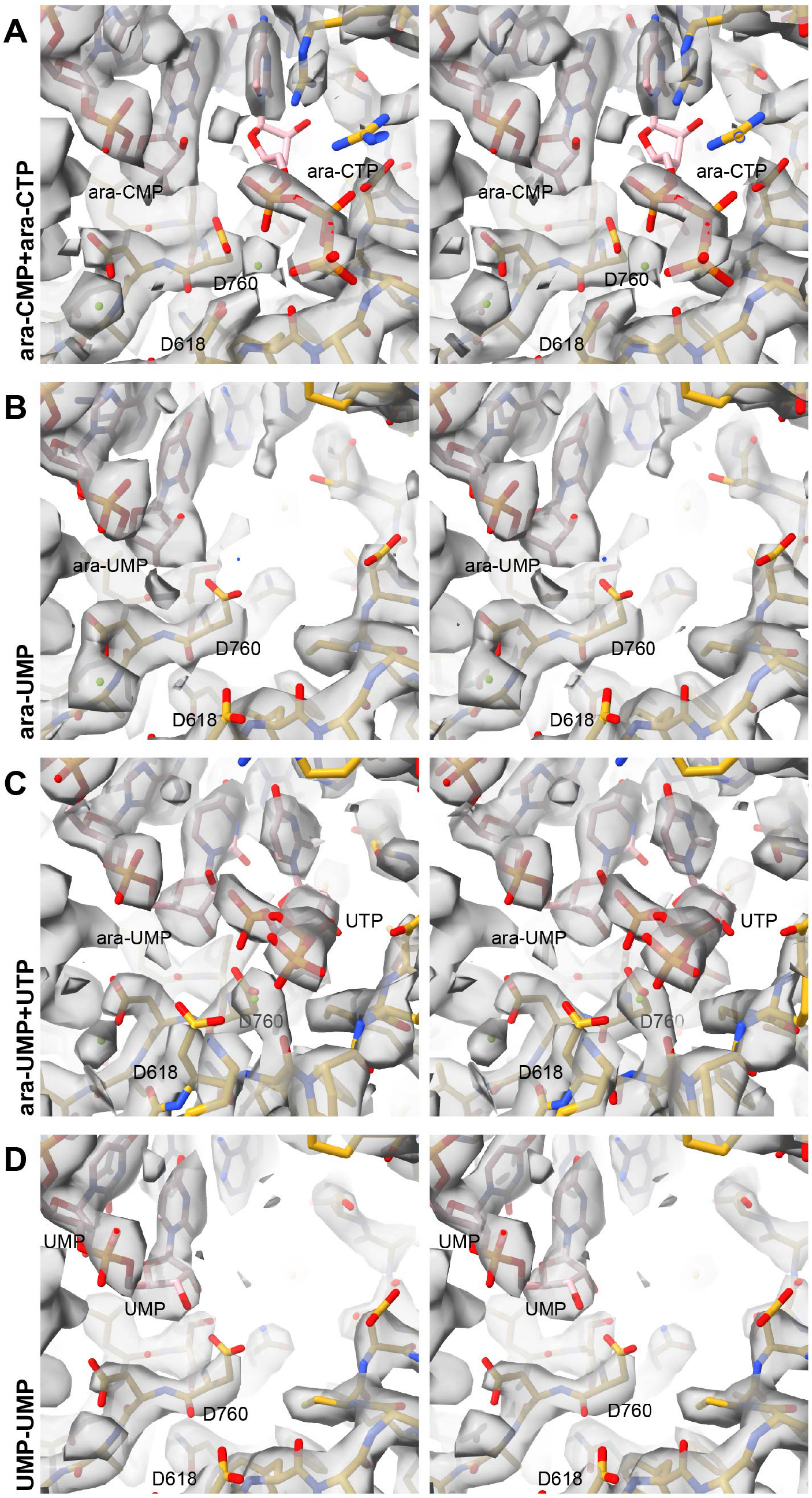


**Figure S8: Stereoimages of determined structures in reconstructed density. A)** Coordinate model (pdb_00009BLF) for the SARS-CoV-2 RTC with incorporated ara-CMP is shown in the corresponding reconstructed density (EMD-44654) isosurface contoured to 0.3. **B)** Coordinate model (pdb_00009PYW) for the SARS-CoV-2 RTC with incorporated ara-UMP is shown in the corresponding reconstructed density (EMD-72038) isosurface contoured to 0.6. **C)** Coordinate model (pdb_00009PYZ) for the SARS-CoV-2 RTC with incorporated ara-UMP and bound UTP is shown in the corresponding reconstructed density (EMD-72053) isosurface contoured to 0.7. **D)** Coordinate model (pdb_00009PZ0) for the SARS-CoV-2 RTC with two incorporated UMP is shown in the corresponding reconstructed density (EMD-72054) isosurface contoured to 0.8.

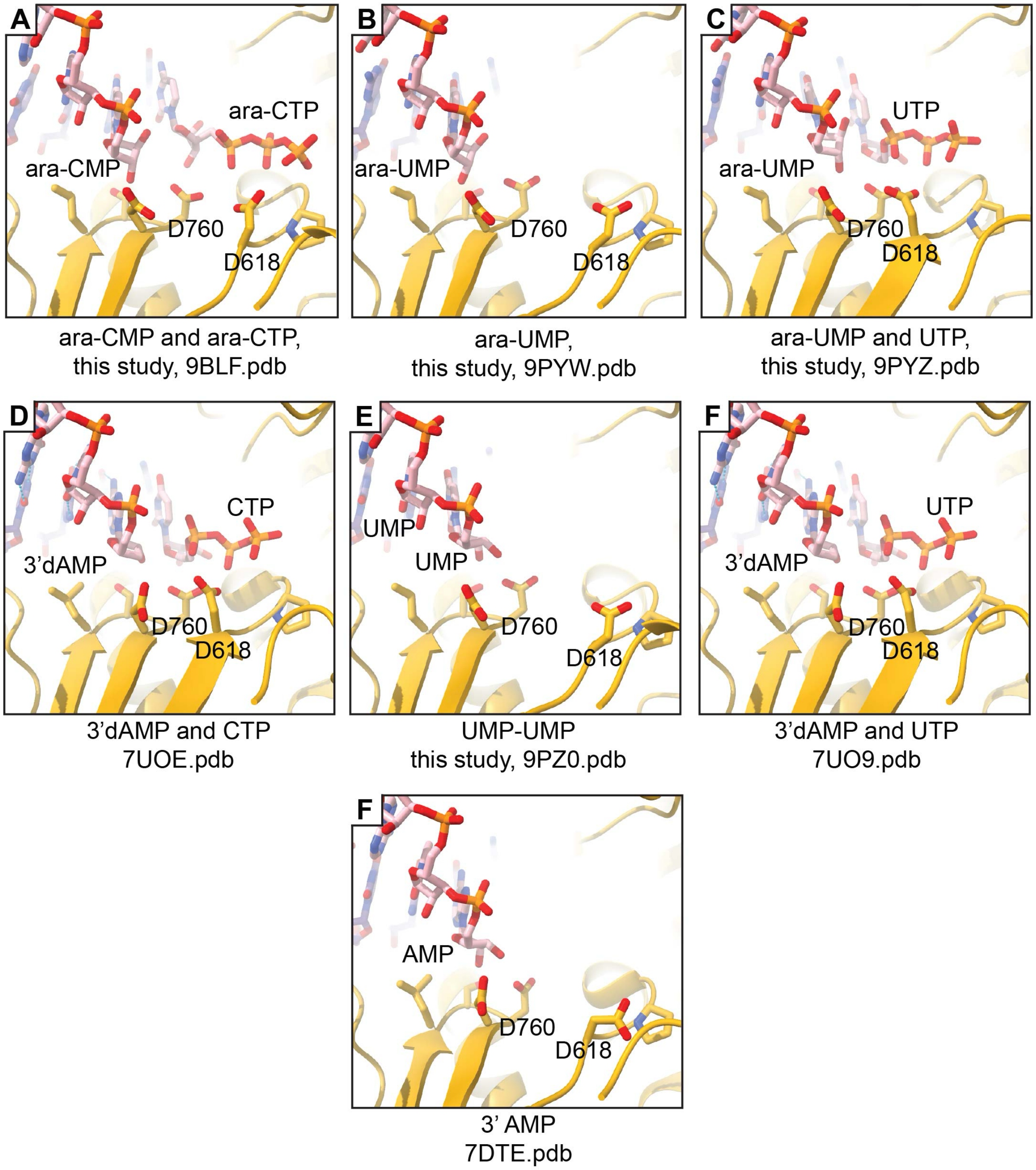


**Fig S9: Comparison of nucleotide bound SARS-CoV-2 polymerase structures.** The figure shows a collection of RNA-bound, SARS-CoV-2 polymerase active sites. Panels **A) B) C)** and **E)** are results from the current study and are described in the text. Panels **D**) and **F)** are derived from Malone et al.[14] who used a 3’ deoxyadenosine to capture natural NTPs in the polymerase active site. Panel **F)** is derived from Wu et al.[16] and depicts and RNA containing a 3’ adenosine.

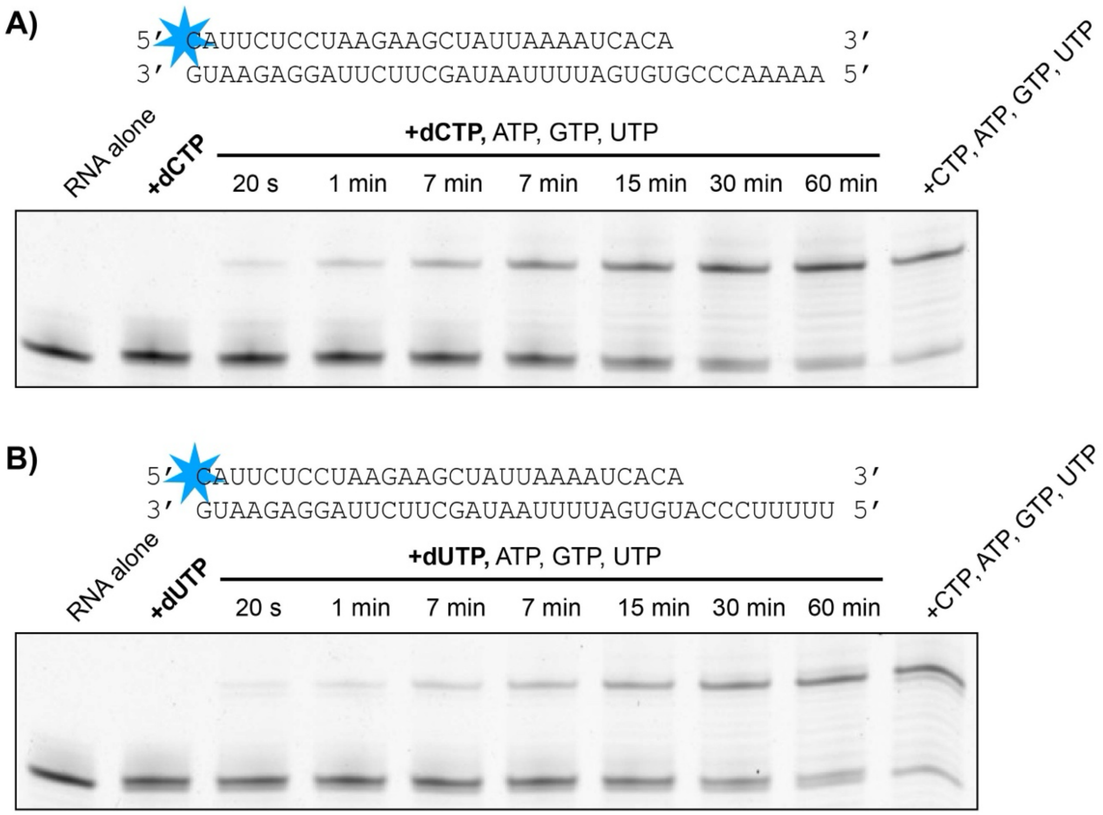


**Figure S10: dNTP Incorporation and chase assay.** Indicated primer template pairs were used to incorporate **A)** dCTP or **B)** dUTP and then the reactions were chased with the remaining nucleotides and followed over time. Primers are labeled with a 5’ 6-FAM tag and reactions are analyzed using denaturing urea-PAGE with FAM detection.

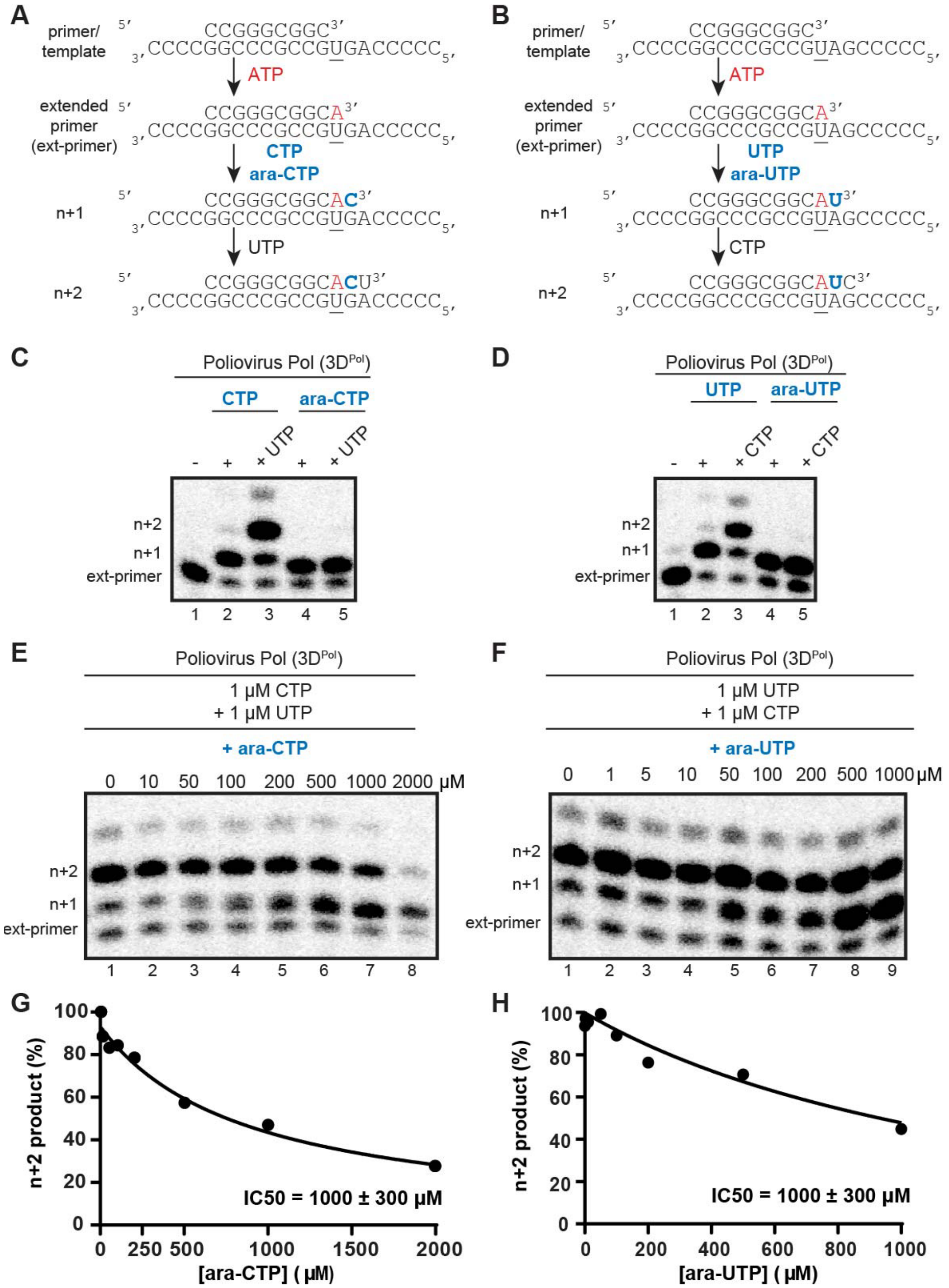


**Figure S11: Incorporation and competition of ara-NTPs. A, B**) Schematic of the primer extension assay used to evaluate poliovirus polymerase $3D^{Pol}$ activity. **C, D**) Incorporation and extension of ara-CTP and ara-UTP. Polymerase-catalyzed incorporation of nucleoside triphosphates using CTP or ara-CTP, and UTP or ara-UTP in the absence and presence of the next correct NTP, respectively. **E-H**) Competition experiment. Polymerase-catalyzed nucleotide incorporation in the presence of increasing concentrations of ara-CTP (0, 10, 50, 100, 200, 500, 1000, and 2000 μM) with 1 μM CTP and 1 μM UTP, and ara-UTP (0, 1, 5, 10, 50, 100, 200, 500, and 1000 μM) with 1 μM UTP and 1 μM CTP. Data was fit to the dose response curve (panels G and H). The $IC_{50}$ values for both ara-CTP and ara-UTP under these conditions are 1000 ± 300 μM.

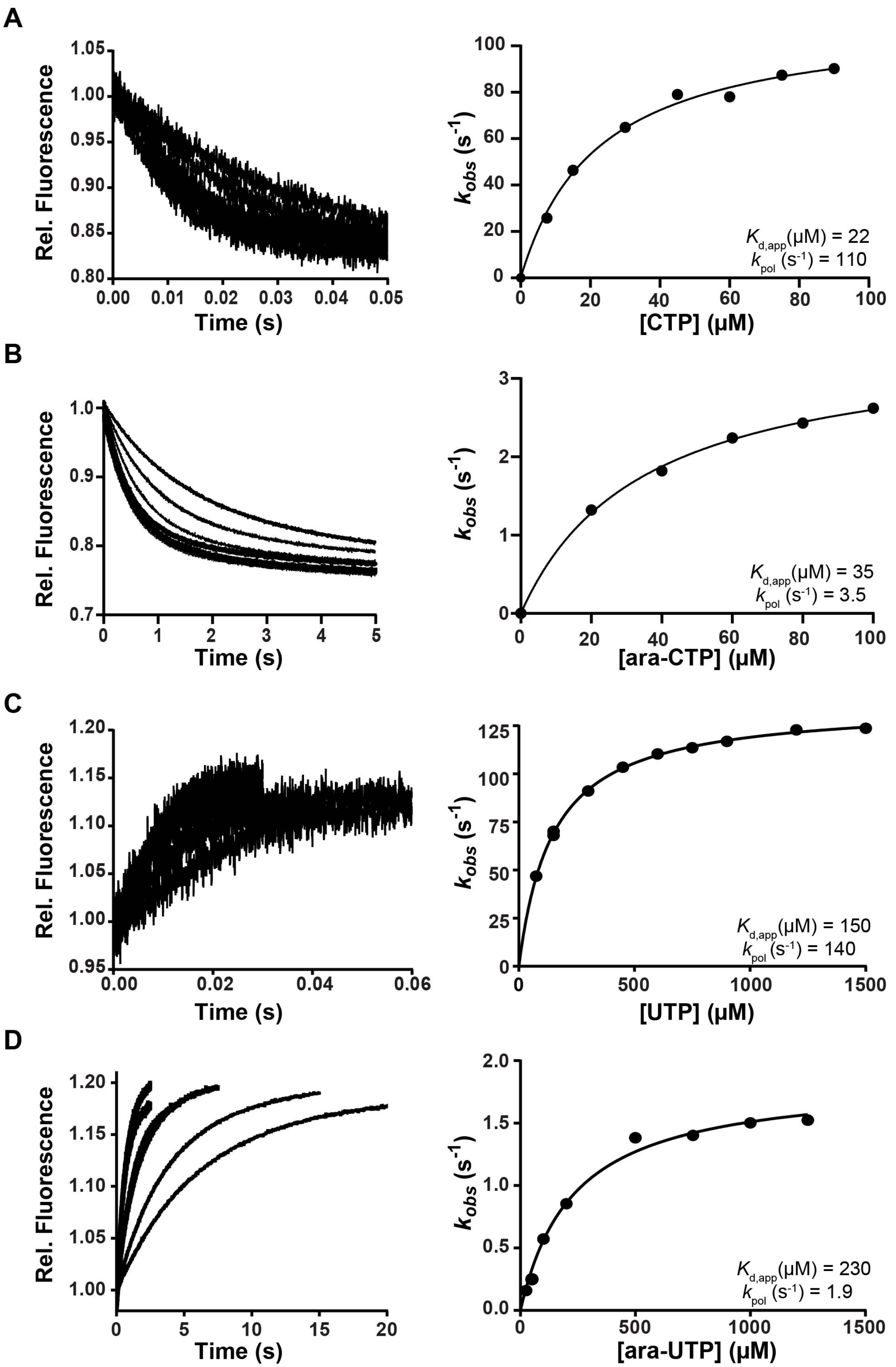


**Figure S12: Kinetics of NTP incorporation.** (**A–D**) Stopped-flow assays were used to determine $K_{d,app}$ and $k_{pol}$ for ara-UTP, ara-CTP, and their natural counterparts. Nucleotide concentrations ranged from 7.5 to 1500 μM, as detailed in the Supplemental Methods. Fluorescence changes were fitted using single-exponential models (UTP, ara-UTP, CTP) or a double-exponential model (ara-CTP) (left panels). The resulting $k_{obs}$ values were plotted against nucleotide concentration (right panels), and fitted to a hyperbolic function to extract $K_{d,app}$ (μM) and $k_{pol}$ ($s^{-1}$).

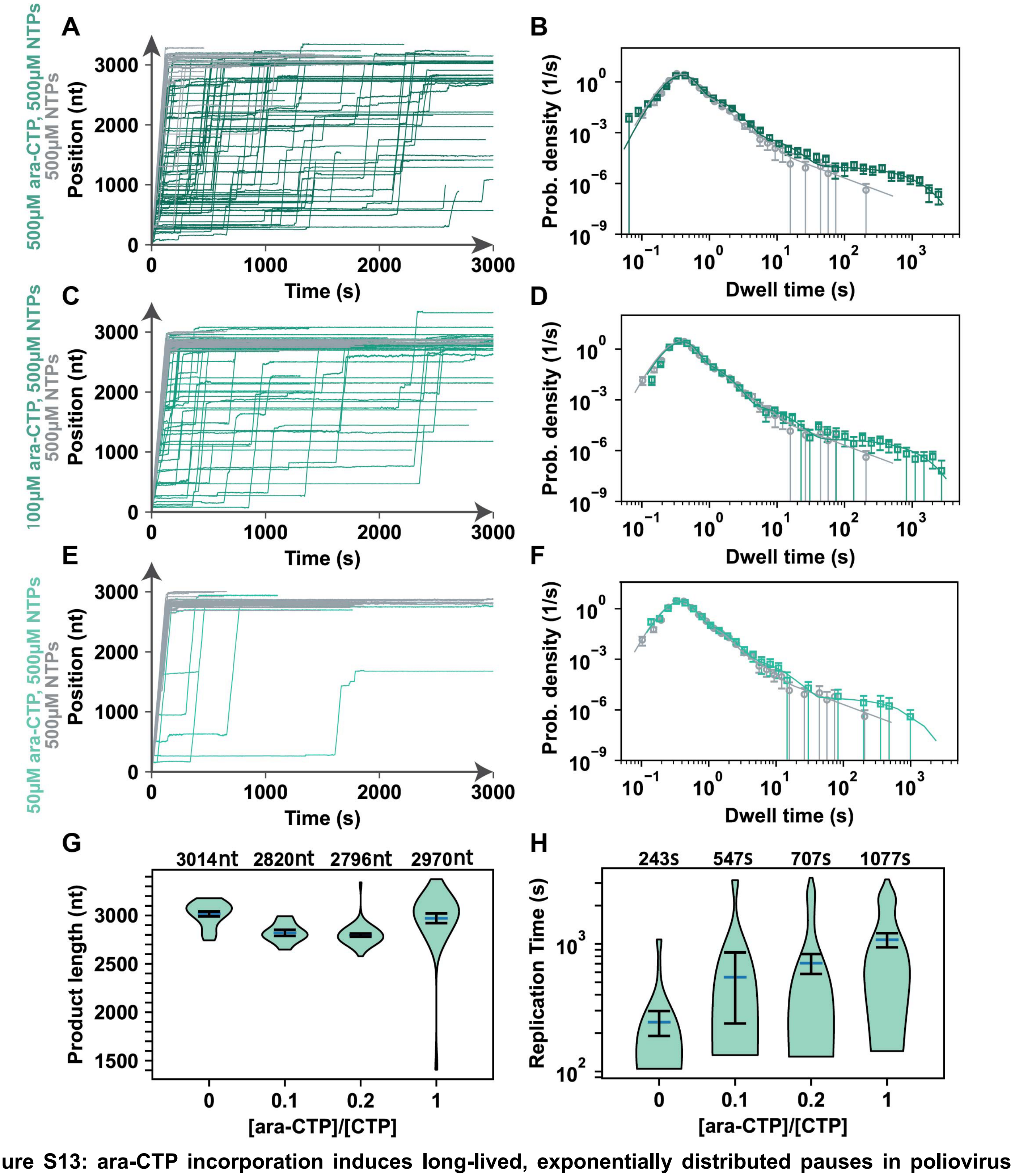


**Figure S13: ara-CTP incorporation induces long-lived, exponentially distributed pauses in poliovirus polymerase elongation dynamics. A, C, E)** Poliovirus polymerase RNA synthesis activity traces comparing 0 µM ara-CTP and 500 µM NTP (grey), or 500 µM, 100 µM, or 50 µM ara-CTP, and 500 µM NTP (green). **B, D, F)** Dwell time distributions extracted from the poliovirus polymerase activity traces in A, C and E respectively. The solid lines are MLE fits of the pause-stochastic model. The error bars are one standard deviation from 1,000 bootstraps of the data sets. **G, H)** Product length and mean replication time of the poliovirus polymerase over 2,820 nt long template as a function of [ara-CTP]:[CTP] stoichiometry. The mean values of the product length and replication time are indicated over the violin plot as dark blue solid line flanked by two horizontal black lines representing one standard deviation error bars extracted from 1,000 bootstraps.

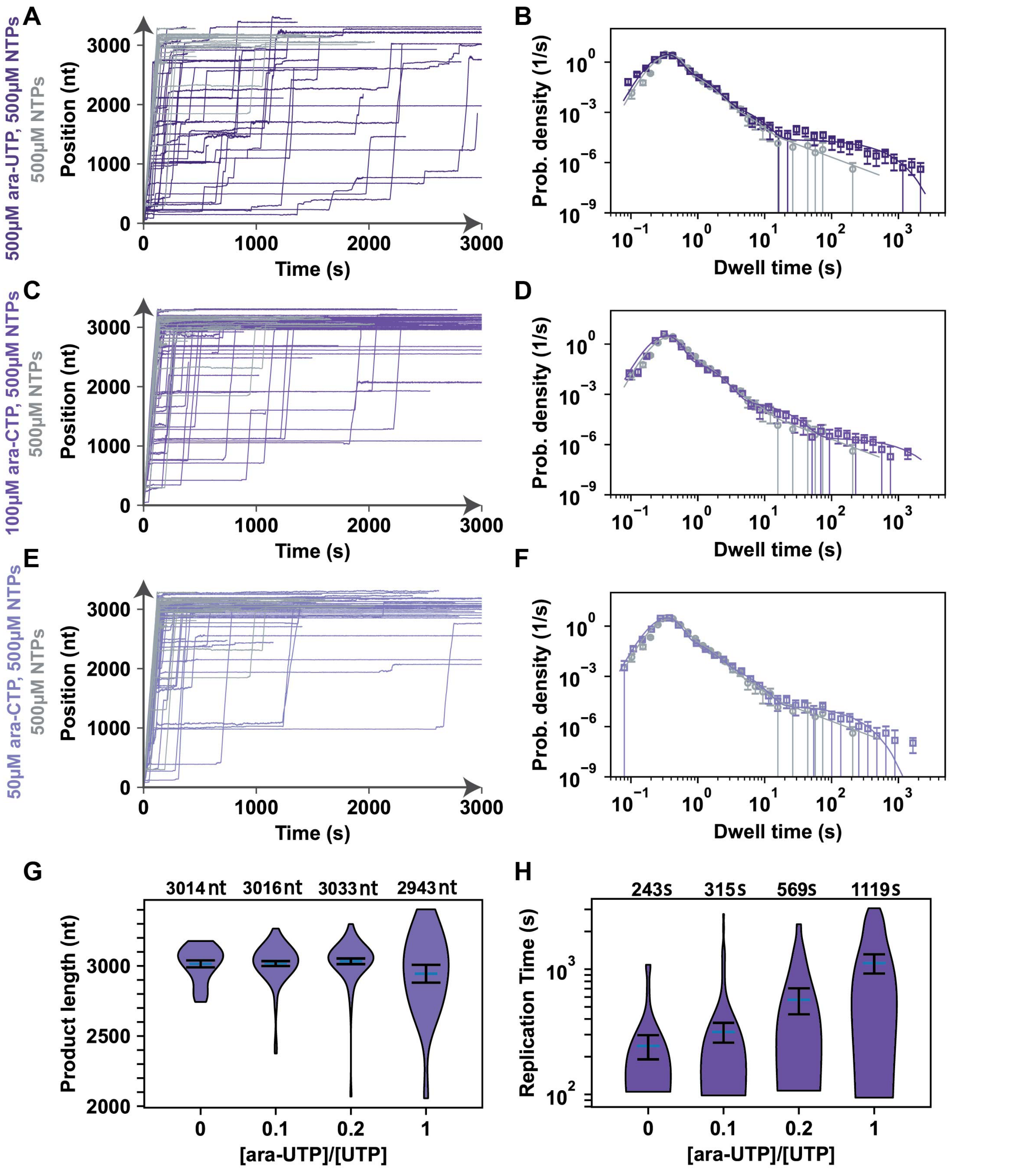


**Figure S14: ara-UTP induces pauses of long duration upon incorporation by poliovirus polymerase. A, C, E)** Poliovirus polymerase RNA synthesis activity traces comparing 0 µM ara-UTP and 500 µM NTP (grey), or 500 µM, 100 µM, or 50 µM ara-UTP, and 500 µM NTP (purple). **B, D, F)** Dwell time distributions extracted from the poliovirus polymerase activity traces in A, C and E respectively. The solid lines are MLE fits of the pause-stochastic model. The error bars are one standard deviation from 1,000 bootstraps of the data sets. **G, H)** Product length and mean replication time of the poliovirus polymerase over 2,820 nt long template as a function of [ara-UTP]:[UTP] stoichiometry. The mean values of the product length and replication time are indicated over the violin plot as dark blue solid line flanked by two horizontal black lines representing one standard deviation error bars extracted from 1,000 bootstraps.

**SUPPLEMENTAL REFERENCES**

Automatic citation updates are disabled. To see the bibliography, click Refresh in the Zotero tab.
